# Characterization of the DNA binding domain of StbA, a key protein of a new type of DNA segregation system

**DOI:** 10.1101/2022.04.29.490116

**Authors:** Valentin Quèbre, Irene del Campo, Ana Cuevas, Patricia Siguier, Jérôme Rech, Phan Thai Nguyen Le, Bao Ton-Hoang, François Cornet, Jean-Yves Bouet, Gabriel Moncalian, Fernando de la Cruz, Catherine Guynet

## Abstract

Low-copy-number plasmids require sophisticated genetic devices to achieve efficient segregation of plasmid copies during cell division. Plasmid R388 uses a unique segregation mechanism, based on StbA, a small multifunctional protein. StbA is the key protein in a segregation system not involving a plasmid-encoded NTPase partner, it regulates the expression of several plasmid operons, and it is the main regulator of plasmid conjugation. The mechanisms by which StbA, together with the centromere-like sequence *stbS*, achieves segregation, is largely uncharacterized. To better understand the molecular basis of R388 segregation, we determined the crystal structure of the conserved N-terminal domain of StbA to 1.9 Å resolution. It folds into an HTH DNA-binding motif, structurally related to that of the PadR subfamily II of transcriptional regulators. StbA is organized in two domains. Its N-terminal domain carries the specific *stbS* DNA binding activity. A truncated version of StbA, deleted of its C-terminal domain, displays only partial activities *in vivo*, indicating that the non-conserved C-terminal domain is required for efficient segregation and subcellular plasmid positioning. The structure of StbA DNA-binding domain also provides some insight into how StbA monomers cooperate to repress transcription by binding to the *stbDR* and to form the segregation complex with *stbS*.

## Introduction

The inheritance of genetic information is a fundamental biological process, essential in all living cells. In bacteria, most chromosomes and low copy-number plasmids are endowed with active segregation (or partition) systems that grant their transmission from parents to offspring. For plasmids, they are composed of three essential components that are necessary and sufficient for their maintenance over generations: a *cis*-acting centromere-like site and two genes, arranged in an operon encoding an NTPase and a centromere-binding protein (CBP) (reviews (1); (2)). The partition process involves: (i) the assembly of a nucleoprotein complex, called the partition complex, by binding of the CBP to the centromere-like site, and (ii) positioning of the partition complexes by the action of the NTPase. This latter involves the separation of the two copies after their duplication, their transport toward opposite cell poles, and their positioning around their new segregated positions until cell division occurs.

Three main groups of partition systems have been described, exemplified by those encoded by plasmids F and P1 (Type I or ParABS), R1 (Type II or ParMRC) and pXO1 (Type III or TubRZC), which are defined by the protein family to which their NTPase protein belongs (2): namely, Walker A/p-loop ATPases, actin-like and tubulin-like, respectively. Type I partition systems are by far the most abundant on low copy-number plasmids and are also the only type encoded by chromosomes. Contrary to the NTPases, the CBP proteins do not show significant sequence similarities, even within the same partition group, although they share structural motifs involved in DNA-binding, in particular their DNA binding motifs. CBPs are dimers composed either of Helix-Turn-Helix motifs (HTH_2_, Type Ia and Type III) or Ribbon-Helix-Helix motifs (RHH_2_, Type Ib and Type II) structural domains (2). Generally, RHH_2_ CBPs cognate centromeres carry arrays of direct repeat sequences that vary extensively in length and organization. In contrast, HTH_2_ CBPs are associated with centromeres containing one or more 13- to 16- bp inverted repeat DNA sequences.

That being said, some low-copy-number plasmids do not encode canonical segregation systems. Among them, two plasmids, the staphylococcal plasmid pSK1 and the enterobacterial plasmid R388, the prototype of the PTU—W family of broad host range conjugative plasmids (3), encode non-canonical systems that involve just a single plasmid-encoded protein devoid of NTPase activity ((4);(5); (1)). These plasmids thus harbor novel segregation systems for which no plasmid-encoded NTPase is present. The partitioning mechanisms of these two unrelated plasmids are not understood.

Plasmid R388 segregation relies on (i) the StbA protein, encoded by the *stbABC* operon, and (ii) on *stbS*, the *cis*-acting centromere-like site located in the promoter of the operon (5). Notably, the *stbABC* operon is close to the origin of transfer (*oriT*) region and is divergently transcribed to the *trwABC* operon involved in conjugative DNA processing (Figure 1). The *stb* operon contains three genes, *stbA*, *stbB* and *stbC*. StbA is the only plasmid-encoded protein required for R388 segregation (5). It was proposed to ensure positioning of plasmid copies in the nucleoid area, since its inactivation leads to aberrant intracellular positioning of plasmid DNA molecules at the cell poles, correlated to plasmid instability without affecting plasmid copy number (Figure 1, (5); (6)). Contrary to what one might expect (by analogy with Par systems) StbB, which contains a putative ATPase motif, is not involved in R388 stability (5). Also, *stbC* encodes a protein with no significant homologs, and its deletion shows no effect in stability and conjugation of plasmid R388 in *E. coli* (5).

**Figure 1.**
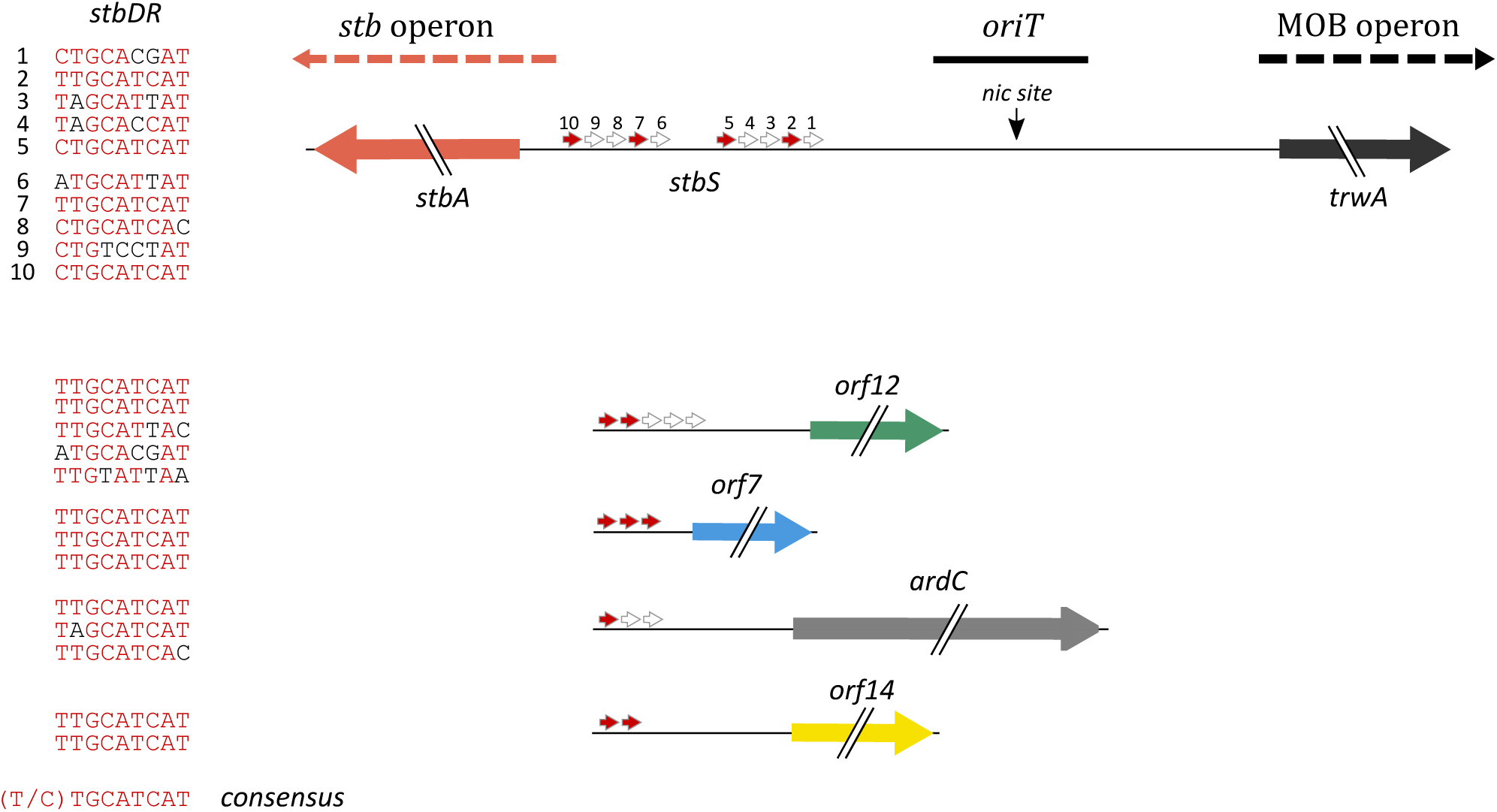
Sequence and positions of the R388 *stbDR* sequences. *stbDR* are represented by gray arrows and consensus ones are colored in red. Genes of which promoters are regulated by StbA and position of *stbS* and the *nic* site of the *oriT* region are indicated. The schematic representation of the *stbDR* sequences and intergenic regions is drawn to scale. DNA sequences of each *stbDR* and the consensus (in red) are shown at the right of the schemas.

In addition to its role in plasmid segregation, StbA acts as a transcriptional repressor of the expression of some R388 genes present in the leading region (*stbA*, *ardC*, *orf7*, *orf12* and *orf14*) (7). All StbA-regulated promoters include two or more direct repeats of a 9-bp DNA sequence with the consensus sequence 5’C/TTGCATCAT, called *stbDR*, separated by 2-bp spacers ((8); Figure 1). The upstream region of the *stb* operon includes the centromere-like site *stbS*, consisting in two sets of five *stbDR. stbS* is strictly required for plasmid R388 stabilization by StbA, but not for conjugation. Besides, StbA, together with StbB, controls the conjugation process but with opposite and interdependent effects. Indeed, while deletion of the entire *stb* operon did not significantly affect conjugation, an activation (50-fold) or a total inhibition of the frequency of conjugation was observed when *stbA* or *stbB* were deleted, respectively. This dual role of StbA hinted for the first time a mechanistic interplay between segregation and conjugation (5) (1). Experimental evidence for functional links between plasmid segregation and conjugation functions have also been described for plasmids R1 and RA3, although by distinct mechanisms (9, 10).

To gain insight into the molecular mechanism by which StbA controls plasmid R388 segregation, we carried out structural, biochemical and *in silico* analyses that characterized the role of the N-terminal domain of the protein. We found that it folds into an HTH motif resembling the DNA-binding domain of the PadR family of transcription factors. We show that the StbA N-terminal domain includes the DNA binding activity required for specific binding to the *stbDR*, as well as for transcriptional repression and for StbA activities in plasmid segregation and in the control of conjugation.

## Materials and methods

### Bacterial strains, plasmids and general procedures

Bacterial strains and plasmids used in this study are listed in Table S1, and oligonucleotides in Table S2. Plasmid pET-StbA_1-75_ was constructed by amplifying the truncated *stbA* gene encoding the first 75 residues of StbA by PCR from plasmid R388 using primers StbAN and StbA75 and introduced between the *Nde*I and *Xho*I restriction sites of pET29c (Novagen). Plasmid R388-StbA_1-75_ was constructed in two steps by replacement of the *stb* operon in R388 (Table S1) by a mutated version carrying a truncated *stbA* gene encoding the 75 first residues of StbA (*stbA_1-75_*). First, a sequence containing the *stb* promoter and *stbA_1-75_* and a sequence containing *stbB* and *stbC* were amplified by PCR from plasmid R388 using primers CG440 and CG387b, and CG441 and CG392, respectively (Table S2). These PCR products were then assembled and cloned into a plasmid pAPT110 linearized with NheI using the In-Fusion® HD cloning kit (Takara Bio) to generate plasmid p4G39. Secondly, R388-StbA_1-75_ was constructed using phage λ red-mediated gene recombination using DY380 strain (11). DNA substrates were generated through PCR amplification from p4G39 with primers CG426 and CG427 that produced a linear DNA fragment containing the mutated *stb* operon and a Kanamycin resistance cassette containing and at least 50 bp terminal arms homologous to the sequences upstream and downstream of the *stb* operon. DNA substrates were introduced by electroporation into DY380 strain harboring R388 grown as described in (12). Cells were plated on agar plates containing Km to select for the desired insertion. Plasmid R388-StbA_1-75_Δ*stbB* was constructed following the same strategy but using primers CG442 and CG443 to generate plasmid p4G41. Constructions were all verified by sequencing of a PCR product of the corresponding region using appropriate primers (see Table S2).

Luria–Bertani (LB) broth was used for bacterial growth (13). For microscopy, M9 medium supplemented with 0.2% casamino acids, 0.4% glucose, 2 μg/ml thiamine, 20 μg/ml leucine and 20 μg/ml thymine was used. Selective media included antibiotics as needed at the following concentrations: ampicillin (Ap), 100 μg/ml; chloramphenicol (Cm), 10 μg/ml; Kanamycin (Km), 25 μg/ml; nalidixic acid (Nx), 20 μg/ml; streptomycin (Sm), 300 μg/ml; spectinomycin (Sp), 30 μg/ml. Plasmid DNA was extracted using PureYield^TM^ plasmid miniprep system (Promega). PCR products were purified using GFX^TM^ PCR DNA and Gel Band Purification Kit (GE Healthcare). Restriction endonucleases and T4 DNA ligase were purchased from Thermo Fisher Scientific and PrimeSTAR DNA polymerase from Takara Bio. Primer oligonucleotides were purchased from Eurofins Genomics.

### In silico analyses

BLASTP searches were performed on the NCBI BLAST online interface (http://www-ncbi-nlm-nih-gov.insb.bib.cnrs.fr/BLAST/) with default parameters. The search for protein domains was carried out using the Interpro (14) and Pfam (15) databases. Multiple alignment was performed using ClustalW (16), the ClustalW algorithm (17) was used for multiple alignments and results were displayed using the Jalview alignment editor (18). The public Galaxy platform at Pasteur was used as an execution engine for web services (19).

### Conjugation and stability assays

Conjugation and stability assays were performed as described in (5). The percentage of plasmids loss (L) per generation were calculated as previously described (20): % L = 1- (Ff/Fi)^(1/n)^x100, where Ff is the fraction of cells carrying the plasmid initially and Fi is the plasmid-carrying fraction after n generations of non-selective growth.

### Protein purification

StbA or StbA_1-75_ proteins were expressed in *E. coli* Rosetta (DE3) cells (Novagen) carrying pET-StbA or pET-StbA_1-75_, respectively. Cells were grown at 37°C to an OD600 nm of approximately 0,6 and protein expression was induced by addition of IPTG to a final concentration of 0.5 mM. After 3 hours growth at 37°C, cells were harvested by centrifugation, resuspended in buffer A (50 mM Tris pH 7.5, 1 M NaCl) and lysed by sonication. Cell-free extracts, obtained by ultracentrifugation at 4°C for 15 minutes at 150 000 g, were loaded onto a 5-ml Ni-affinity column (HisTrap HP, GE healthcare) and washed with buffer B (50 mM Tris pH 7.5, 0.5 M NaCl) containing 5 mM and 50mM Imidazole. Protein was eluted with a 0.05 to 0.5M Imidazole gradient. Fractions containing the protein were pooled, dialyzed against buffer C (20 mM Tris pH 7.5, 0.2 M NaCl, 1 mM EDTA) overnight at 4 °C, and then loaded onto a Superdex S75 gel-filtration column (GE healthcare) and eluted with buffer C. Fractions containing the protein were pooled and either mixed with cold glycerol (20 % final) and DTT (5 mM) and kept at – 80°C or used directly for crystallization trials.

Expression and purification of Selenomethionine substituted StbA_1-75_ was performed as described above with the following modifications: pET- StbA_1-75_ was introduced in B834 (DE3) *E. coli* cells (Novagen) and the resulting strain was grown at 37°C in 1 liter of SelenoMet^TM^ Medium Base (Molecular Dimension) supplemented with selenomethionine and all of the natural amino acids excluding methionine.

### Structure determination (crystallization, data collection and structure determination)

Screening for crystallization conditions was performed using commercially available kits (Hampton Research). Crystals were grown at 19°C using the sitting-drop vapor diffusion technique (1 µl protein solution and 1 µl crystallization reagent equilibrated against a 0.5 ml reservoir volume) upon mixing the protein 1:1 with well solution containing 0.1 M Trisodium Citrate Dihydrate pH5.6, 10% (v/v) iso-propanol and 10% (v/v) PEG 4000. Datasets were obtained at beamline PROXIMA at the SOLEIL Synchrotron Radiation Facility (Gif-Sur-Yvette, France).

For data collection, StbASeMet crystals were flash-frozen in liquid nitrogen at 105 K. For single StbA-SeMet crystals, data was collected at 0.9793Å, the wavelength corresponding to the Selenium absorption maximum according to the fluorescence scan. Diffraction images were processed using XDS (21) and scaled using Scala (22) as part of the CCP4 package (23). The structure was solved by single anomalous dispersion (SAD) phasing using the program AutoSol of the PHENIX package (24). The refinement of the initial model was performed through several cycles by Phenix refine (24) until appropriate R factors were reached. Final manual modeling was done in COOT (25). The depiction of structures and analyses were performed with UCSF Chimera, developed by the Resource for Biocomputing, Visualization, and Informatics at the University of California, San Francisco, with support from NIH P41-GM103311.

### Limited proteolysis

StbA at a concentration of 1.2 mg.ml^-1^ was incubated with several amounts of trypsin (Thermo Fisher Scientific) for different times at 37°C. After trypsin treatment, protein loading buffer (2X: 0.5 M Tris-HCl [pH 6.8], 4.4% [wt/vol] SDS, 20% [vol/vol] glycerol, 2% [vol/vol] 2-mercaptoethanol, and bromophenol blue) was added to stop the reaction. Samples were boiled for 5 min and loaded on a 15% SDS-PAGE. The band corresponding to the major cleavage product was excised for mass spectroscopy identification.

### Electrophoretic mobility shift assays (EMSA)

Fluorescent DNA substrates were prepared by hybridization of complementary Cy3- or Cy5-labeled oligonucleotides (Table S2). Fluorescently labeled probes were incubated in 0.5 M Tris-HCl pH7.5 and 0.1 M NaCl at 95°C for 10 min, and slowly cooled down to 25°C. 67 nM of fluorescent DNA substrate was incubated with increasing concentrations of StbA or StbA1-75 (0 to 16 μM) in a final volume of 12 μl of binding mixture (10mM Tris-HCl (pH7.5), 200 mM NaCl, 0.5 mM EDTA and 20% glycerol) with sonicated salmon sperm DNA as a competitor (100 μg.ml^-1^) for 20 min at 30°C. Reaction mixtures were separated on a 5% non-denaturing polyacrylamide gel in TGE buffer (25 mM Tris, 25 mM Glycine and 5 mM EDTA) and analyzed using a Typhoon trio imager (GE Healthcare). For the short–Long EMSA Assay Coupled to Differential Fluorescent DNA Labeling, the protocol is as described in (26).

### Transcriptional regulation activity measurements

Reporter plasmids carrying *P*stbA, *P*orf7, *P*orf12, *P*orf14 promoters were constructed in a previous study (7). *stbA* and *stbA_1-75_* genes (encoding the first 75 residues of StbA) were amplified from R388 by PCR with primers StbAsen and StbAasen StbA_1-75_sen and StbA_1- 75_asen, respectively, (Table S2) and cloned in plasmid pBAD33 using *Xba*I and *Hind*III restriction endonucleases to generate pBAD*stbA_1-75_* (Table S1). To determine the effect of StbA and StbA_1-75_, plasmids pBAD*stbA* and pBAD*stbA_1-75_* were transformed to *E. coli* Bw27783 containing one of the reporter plasmids. Protein expression was induced by adding appropriate concentrations of arabinose to M9-broth and fluorescence per OD unit (GFP/OD) was determined and compared to that produced by the same reporter strain when containing the empty expression vector pBAD33.

### Microscopy

Live cell microscopy experiments were performed as described in (5) with the following modifications. To fluorescently label bacterial nucleoid, *E. coli* strain LN2666 was modified by P1 transduction to carry a *hupA∷mcherry* translational fusion that expresses the nucleoid associated protein HU fused with mCherry (Table S1; (27). Strain LN2666 Hu-mcherry containing plasmid pALA2705 (Table S1) was transformed with DNA of plasmid R388::*parS-Cm* (R388) or one of its derivatives. Cells were visualized at 30°C using an Eclipse TIE/B wide field epifluorescence microscope (Nikon) with a phase contrast objective (CFI Plan Apo Lbda 100X oil NA1.45) and a Semrock filter FITC (Ex: 482BP35; DM: 506; Em: 536BP40) or mCherry (Ex: 562BP40; DM:593; Em: 641BP75).

Images were acquired using an Andor Neo SCC-02124 camera with illumination at 60% (FITC) or 25% (mCherry) from a SpectraX source Led (Lumencor) and exposure times of 800 and 100ms respectively for FITC and mCherry. Nis-Elements AR software (Nikon) was used for image capture and editing. Foci detection and integrated fluorescence phase-contrast and fluorescence were measured using Coli-Inspector project running under ImageJ software in combination with plugin ObjectJ (http://simon.bio.uva.nl/objectj/). At least 1000 cells were inspected for each experiment.

## Results

### StbA is organized in two domains

Comparative genomic studies previously showed that *stbA* is located in a region of conserved synteny, adjacent to the origin of transfer of plasmids of several mobility groups (MOB_F11_, MOB_P11_, MOB_P6_, MOB_P13/P14_) belonging to various PTUs (W, N1, P1, P9, Q1 and I2) ((5); (3)). In order to gain structural information and assess the diversity of the StbA proteins, we performed BLASTP searches among all complete prokaryotic genome sequences available (BLAST+2.11.00) using R388 StbA (Genebank BAD24117) as a query. We analyzed the genetic neighborhood of the StbA homologs and retained those for which the synteny of the *stb* and MOB operons was conserved. Results were then filtered both for redundancy and for keeping a subset of representatives of StbA diversity (see Materials and methods). Figure 2 displays a multiple sequence alignment of R388 StbA and 33 other selected StbA protein sequences. StbA proteins are small proteins of about 150 amino acid residues, ranging from 135 to 175. Analyses from StbA sequences by using the InterPro and Pfam databases did not reveal any known domain, motif or protein family except a disordered region encompassing residues 69 to 108. However, protein sequence comparison showed that the N-terminal half of StbA displays high degree of conservation (Figure 2A), including several blocks of conserved residues, while the C-terminal half is poorly conserved (Figure 2B). These data thus suggest that the StbA proteins are organized in two domains.

**Figure 2.**
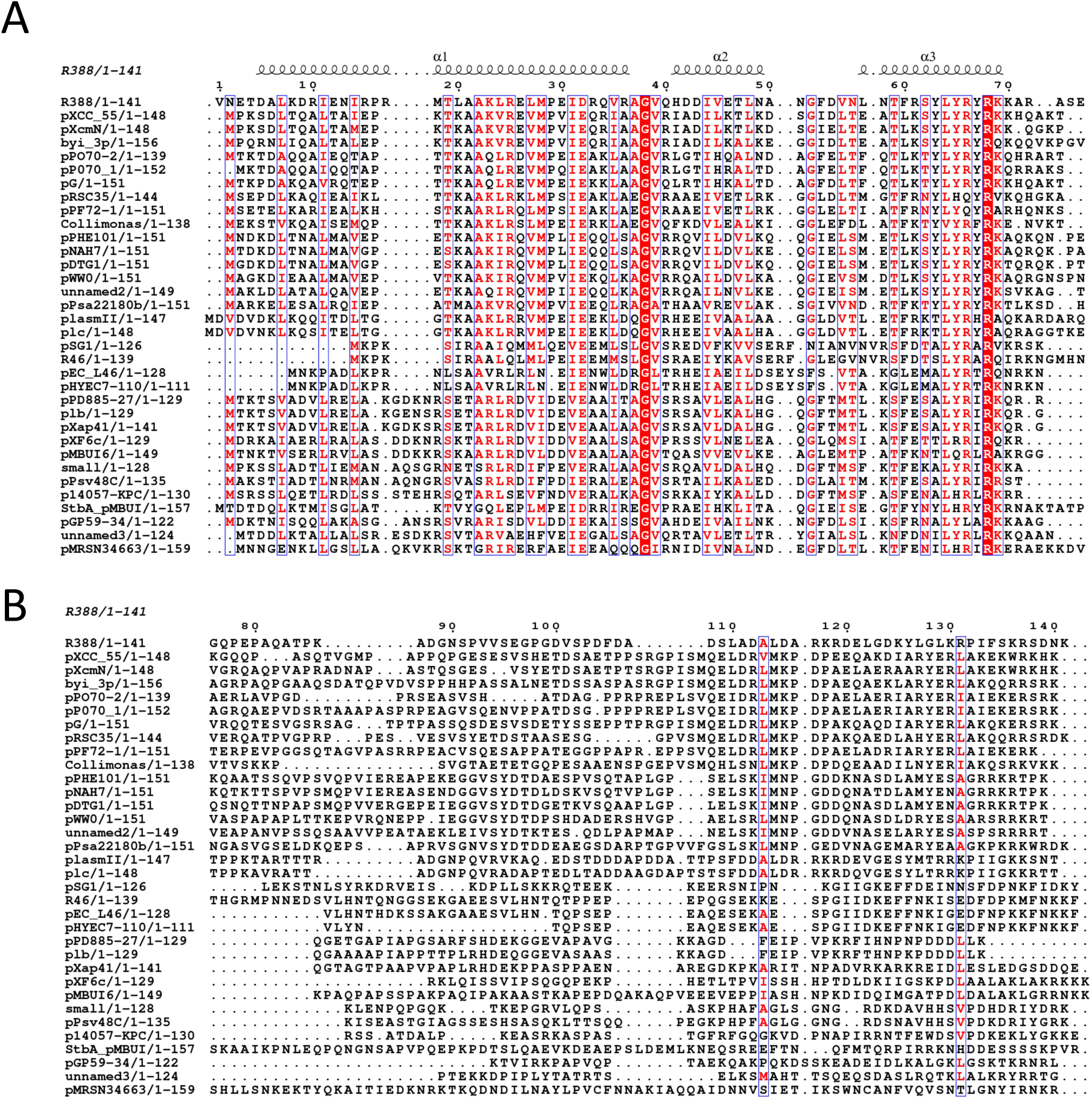
Multiple sequence alignment of StbA protein sequences. The amino acids sequence of StbA from plasmid R388 was used as a query in a BLASTP search among all complete prokaryotic genome sequences available (December 2020). The multiple sequence alignment is adorned by secondary structures elements observed in StbA_1-75_ crystal structure. The figure has been prepared using the Espript 3.0 web server (60). **A**: N-terminal domain, **B**: C-terminal domain.

To experimentally probe the domain organization of R388 StbA, a C-terminally (His)_6_-tagged version of full-length StbA (16,7 kDa) was subjected to time-limited proteolysis with trypsin and analyzed by SDS-PAGE. As shown in figure 3, StbA digestion yielded several discreet proteolytic products (lanes 4-6 and 1-2). The smallest polypeptide band (P) was the most resistant to proteolysis (lane 2), prior to be fully degraded after ∼ 30 min (lane 3). MALDI-TOF spectrometry analysis revealed a fragment of about 10 kDa that corresponds to residues 1 to 86. No peptide corresponding to the last 55 C-terminal residues of StbA was detected in the mass analysis, indicating that the C-terminal part of StbA is a poorly structured region and thus rapidly degraded.

**Figure 3.**
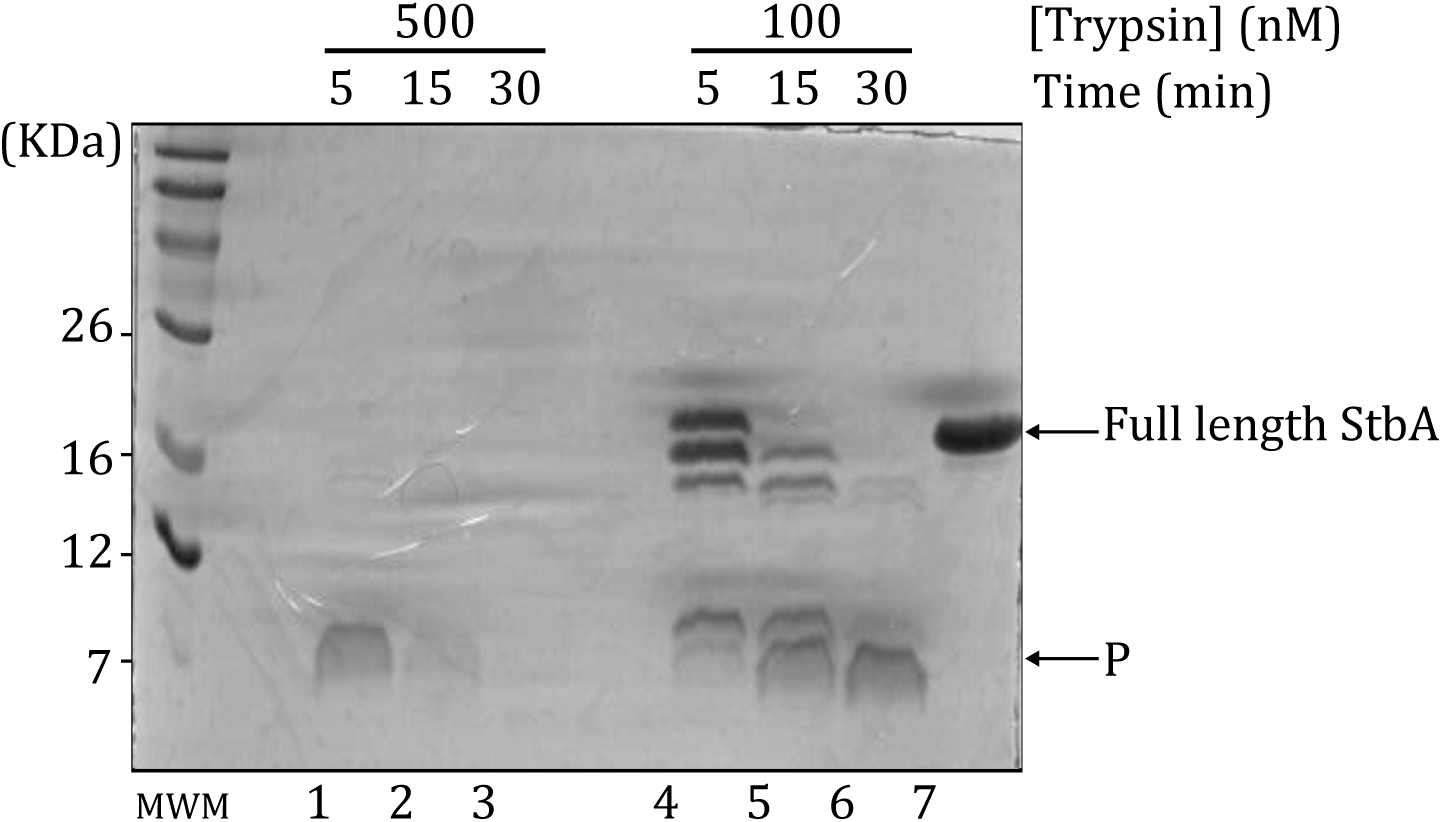
Limited proteolysis of StbA. Purified StbA protein was incubated with different concentrations of trypsin. Samples were taken at different time points (from 5 to 30 minutes) and analyzed by SDS-PAGE on a 10% gel. Molecular weight (in kDa) of the size markers (MW), full-length StbA_His6_ and the proteolytic fragment analyzed by mass spectroscopy (P) are reported on the sides of the gel. Lanes 1-3: proteolysis of purified StbA with trypsin ratio enzyme:protein of 1:1 (w/w). Lanes 4-6: proteolysis of purified StbA with trypsin ratio enzyme:protein of 1:3 (w/w).

### Crystal structure of the N-terminal domain of StbA

For a further characterization, we performed structural studies of StbA. The full length StbA protein was refractory to crystallization, possibly due to its unstructured C-terminal domain. Thus, we overexpressed and purified the N-terminal domain of StbA, spanning residues 1 to 75 with a C-terminal His-Tag (StbA_1-75_-HT).

We solved the crystal structure of StbA_1-75_-HT at 1.9 Å resolution by single anomalous dispersion (SAD) using a selenomethionine-derivative protein (Materials and Methods). The crystal belonged to the space group P3_1_ with one molecule per asymmetric unit. The final model of StbA_1-75_ contains 72 residues (Asn_2_ to Ala_73_) and 12 water molecules. Data collection and refinement statistics are given in Table 1.

**Table 1:** Data collection and model refinement statistics.

|  |  |  |  |
| --- | --- | --- | --- |
| **Wavelength** | 0.9793 | **Reflections used for R-free** | 312 (30) |
| **Resolution range** | 34.4-1.9 (2.0-1.9) | **R-work** | 0.2003 (0.2935) |
| **Space group** | P 3<sub>1</sub> | **R-free** | 0.2222 (0.2688) |
| **Unit cell** | a=b=45.62 c=34.39 | **CC(work)** | 0.954 (0.764) |
| $\alpha=\beta=90$ $\gamma=120$ | | **CC(free)** | 0.928 (0.722) |
| **Total reflections** | 35568 (4758) | **Number of non-H atoms** | 608 |
| **Unique reflections** | 6294 (897) | **Protein residues** | 596 |
| **Multiplicity** | 5.7 (5.3) | **RMS(bonds)** | 0.008 |
| **Completeness (%)** | 99.3 (96.8) | **RMS(angles)** | 0.92 |
| **Mean I/sigma(I)** | 28.5 (3.8) | **Ramachandran favored (%)** | 98.57 |
| **Wilson B-factor (Å<sup>2</sup>)** | 32.05 | **Ramachandran allowed (%)** | 1.43 |
| **R-merge** | 0.032 (0.386) | **Ramachandran outliers (%)** | 0.0 |
| **CC1/2** | 0.999 (0.864) | **Rotamer outliers (%)** | 3.2 |
| **CC\*** | 1.0 (0.907) | **Clashscore** | 5.01 |
| **Reflections used** | 6276 (622) | **Average B-factor** | 34.91 |
| **in refinement** |  |  |  |

The overall architecture of StbA_1-75_ is shown in Figure 4A. It shows a simple fold comprising three α-helical segments with a long N-terminal helix α1 (residues 5–36), followed by two short helices α2 (residues 41–49) and α3 (residues 57–69). α2 and α3 helices are connected by a sharp turn of 7 residues (residues 51-56, Figure 4A). This fold corresponds to the characteristic triangular outline that defines the tri-helical HTH domain (for reviews see (28) and (29)). HTH motifs are well-known DNA-binding domains present in the three super-kingdoms of life. They are found in the most prevalent transcription regulators of all prokaryotic genomes and are also involved in various other functions, such as DNA repair, replication and RNA metabolism and also in plasmid partition (30).

**Figure 4.**
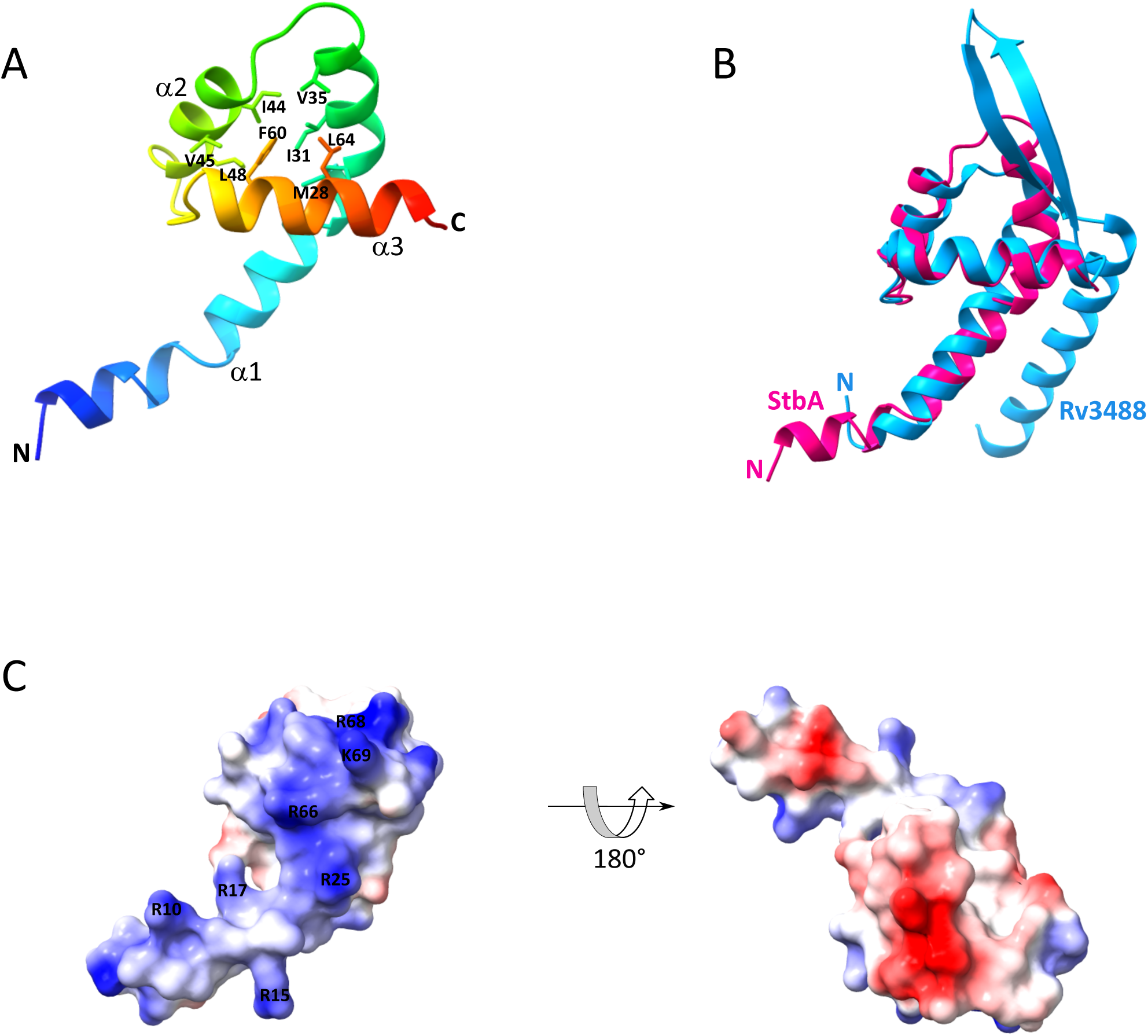
Crystal structure of StbA N-terminal domain. **A.** Ribbon diagram of the monomeric tri-helical HTH structure of StbA_1-75_ showing the HTH fold formed by the three helices in the molecule. Secondary structure elements and conserved residues forming the hydrophobic core of the HTH motif are labelled. **B.** Structural superposition of StbA_1-75_ (crimson) with the PadR-like transcriptional regulator Rv3488 of *Mycobacterium tuberculosis* H37Rv (blue, PDB ID: 5ZHC). **C.** Electrostatic surface potentials of StbA_1-75_ showing a conserved positive surface patch. Positive, negative, and neutral electrostatic potentials are represented by blue, red, and white, respectively. Most conserved basic residues are shown.

StbA HTH motif displays conserved sequence elements that are characteristic of the HTH fold. The sharp turn between α2 and α3, which is a defining domain of this fold, is very conserved in the StbA proteins (Figures 2 and S1). It contains a motif Gly_52_-Phe_53_-Asp_54_, which corresponds to a ‘Gha’ pattern (where G is a glycine, h a hydrophobic residue, and an acidic residue), reminiscent of the characteristic ‘shs’ pattern (where s is a small residue; (29); Figure S1). Also, other conserved hydrophobic residues point to the interior of the fold, forming a characteristic hydrophobic core that stabilizes the domain (residues Met_28_, Ile_31_ and Val_35_ in α1, Ile_44_, Val_45_ and Leu_48_ in α2, and Phe_60_ and Leu_64_ in α3, Figure 4A).

A Dali structural similarity search indicated that StbA_1-75_ structure most closely resembles the winged HTH motif of the transcriptional regulator Rv3488 of *Mycobacterium tuberculosis* H37Rv, which belongs to the PadR-like family (Figure 4B; PDB ID: 5ZHC; (31) with a Dali Z-score of 6.4 and an r.m.s. deviation of 2.5 Å (with an overall sequence identity of 17%).

PadR-like transcriptional regulators form a large structurally-related family of proteins that play roles in diverse biological processes, such as detoxification, virulence, antibiotic synthesis and multi-drug resistance in various bacterial phyla ((32); (33); (34); (35); (36); (37)). The AspA CBP of the archeal *sulfolobus* plasmid pNOB8 partitioning system also shows similarity with the PadR proteins (38). The structural similarity between StbA_1-75_ and PadR-like DNA-binding motif indicates that StbA contains a functional HTH motif.

PadR-like regulators share a common fold comprising a highly conserved N-terminal winged-HTH domain, consisting of three α helices (α1 to α3), a two C-terminal β-strand hairpin unit (the wing), and a variable C-terminal module involved in dimerization. StbA_1-75_ lacks the wing, which often provides an additional interface for DNA contact through charged residues in the hairpin (29). Although full-length StbA eluted as a dimer in exclusion chromatography, StbA_1-75_ appears to be a monomer in solution (Figure S2). This is consistent with crystallographic data showing that the asymmetric unit (a.s.u.) contained one StbA_1-75_ monomer. On the other hand, our analyses of StbA_1-75_ using the bacterial adenylate cyclase-based bacterial two-hybrid system showed that it self-associated, indicating that the protein forms multimers *in vivo* (BACTH, Figure S3). Although these observations could result from crystallization artifacts, the crystal structure shows that the N-terminal part of helix α1 of one monomer packs into a hydrophobic pocket formed between helices α1 and α2 of another monomer. Positions of the hydrophobic residues putatively involved in interactions at the dimer interface, which are mostly conserved in StbA proteins, are indicated in Figure S4 (Leu_20_, Leu_27_, Ile_31_, Val_39_, Ile_44_ and Ile_48_ from monomer 1, and Leu_7_, Ile_11_ and Ile_14_ from monomer 2). The crystals showed a three-fold symmetry between three identical subunits. Although this observation may be an artifact due to crystallization and/or the absence of the C-terminal domain of StbA, it suggests that StbA_1-75_ monomers may interact as trimers. In the structure, a hydrogen bond formed by Asp_5_ from monomer 1 and Lys_69_ from monomer 3 may stabilize the interactions between dimers (Figure S4B). Oligomerization of StbA may also involve the C-terminal domain of the protein, as described for several PadR-like proteins ((38); (39); (36); (31); (40)).

The third helix, α3, is known as the recognition helix that inserts into the major groove of the DNA to form the primary protein-DNA interface (41). As shown in Figure 2, StbA α3 contains mainly conserved polar residues (Asn_56_, Thr_59_, Ser_62_), including several positively charged residues (Arg_61_, Arg_66_, Arg_68_ and Lys_69_), which are presumably involved in specific protein-DNA interactions. The electrostatic model presented in Figure 4C shows that StbA_1-75_ displays a large highly positively charged surface extending over the whole side of the protein exposing α3, which contains many basic amino residues. These are contributed from α3 (see above) and also the N-terminus of α1 (Arg_10_, Arg_15_, Arg_17_ and Arg_25_), which suggests that α1 could be involved in additional protein-DNA contacts with the minor groove of the DNA. Consistently, Arg_25_ and Arg_17_ are pretty well conserved (Figure 2). This putative DNA attachment site involving the N-terminus of α1 is characteristic of the HTH motif of the homeodomain family (28). Notably, the structure of StbA N-terminal domain resembles the homeodomain fold ENT of the eukaryotic protein EMSY (Z-score 5.0, rmsd=4.9) (42).

### StbA binding on the *stbDRs*

StbA was shown *in vitro* to bind specifically to a 200-bp DNA substrate containing the two sets of five *stbDR* representing *stbS* (5). To further analyze the DNA binding properties of StbA to the *stbDR* sites, we tested by EMSA its ability to bind different numbers of the *stbDR* consensus sequence 5’-T/CTGCATCAT on 80-bp DNA fragments (see figure 1) in the presence of an excess of non-specific competitor DNA (Figure 5).

**Figure 5.**
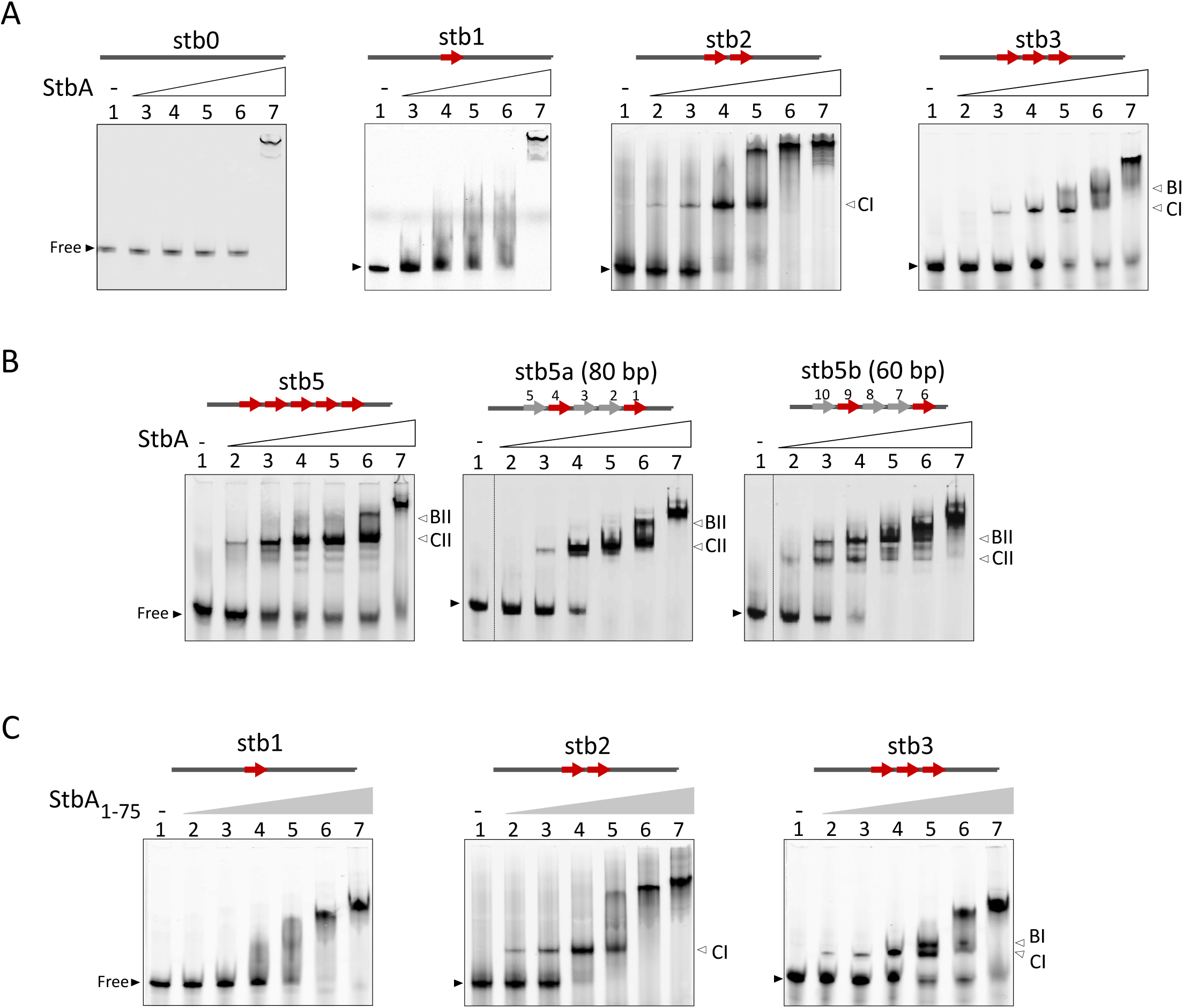
DNA-binding activities of StbA and StbA_1-75_ by EMSA. Fluorescently labeled DNA fragments carrying a variable number of *stbDR* were incubated with increasing concentrations of purified His-tagged StbA **(A, B)** or StbA_1-75_ **(C)**. EMSA were performed on 5% polyacrylamide gels with 0 (lanes 1), 500 nM (lanes 2), 1 μM (lanes 3), 2 μM (lanes 4), 4 μM (lanes 5), 8 μM (lanes 6), and 16 μM (lanes 7) StbA (**A and B**) or StbA_1-75_ (**C**). DNA substrates are schematized at the top of the gels. Arrows represent *stbDR* sequences with the same color legend as in Figure 1. stb1, stb2, stb2b, stb3 and stb5b are 5’-Cy3-labeled, and stb5 and stb5a are 5’-Cy5-labeled. stb5a and stb5b contain *stbDR* 1 to 5 and *stbDR* 6 to 10, respectively. All substrates are 80 bp long except stb5b (60 bp). Dotted lines between lanes indicate that they were not side-by-side in the original gel. Free DNA and protein-DNA complexes are indicated by black and white wedges, respectively, and larger complexes termed high molecular weight complexes (HMW) are shown by vertical bars.

Incubation of increasing amounts of StbA with the DNA substrate carrying a single *stbDR* (stb1, Figure 5A), but not with none (stb0, Figure 5A), gave rise to a smear, indicating specific but unstable interactions (lanes 3-6). Band shifts observed at the highest concentration of StbA (16 μM; Figure 5A, stb1, lanes 7) were considered as non-specific binding since they were also present with the stb0 DNA substrate (Figure 5A, stb0, lane 7). Then, complexes formed at this highest concentration of StbA were difficult to interpret since they probably include both non-specific complexes and specific complexes (lanes 7).

With the DNA substrate carrying 2 *stbDR* (separated by 2 bp as observed in the *StbDR* region), a discrete complex was readily observed (CI, stb2, Figure 5A, lanes 2-5), suggesting that StbA binds to two contiguous *stbDR* sites with cooperativity. However, the substrate including 2 *stbDR* separated by a longer spacer (43 bp) (stb2’, Figure S5A) gave rise to a smear, i.e. a similar pattern as observed with stb1, showing that 2 *stbDR* sequences close to each other are required to form stable complexes (stb2, lanes 2-5). Addition of a third *stbDR* site gave rise to another band (BI) migrating slower than CI (BI, stb3, lanes 5-6). The smear between BI and CI, and the remaining of free substrate at high StbA concentrations suggested that the interactions between StbA and the DNA substrates carrying 3 *stbDR* are less stable than with 2 *stbDR* (compare stb2 and stb3, lanes 4-6). We next examined the binding of StbA to a DNA substrate carrying 5 consensus *stbDR* (stb5, figure 5B). Two major complexes were observed: the major discreet one (CII) displayed an electrophoretic mobility lower than CI, and the second one migrating slower (BII, left panel, lanes 2-6). These data suggest that StbA binds with high cooperativity as a dimer to 2 *StbDR* (CI), and two dimers to 4 *StbDR* (CII). According to this hypothesis, BI and BII could therefore correspond to less stable complexes including an additional dimer bound to the remaining *StbDR* in stb3 and stb5, respectively.

We next tested two other probes carrying 5 *stbDR* but with the 5 *stbDRs* included in the first or the second array of *stbS*, stb5a and stb5b, respectively. Stb5a and Stb5b substrates gave rise to a similar pattern to that observed with stb5 with two major complexes that might correspond to CII and BII (Figure 5B, central panel). With stb5b, which is smaller (60 bp), one more major complex migrating faster is observed (lanes 2-6, Figure 5B, right panel). Notably, free unbound substrate was observed at high StbA concentrations (lanes 5-7) only with stb5, which carries five consensus *stbDR*. These results suggest that the three substrates carrying 5 *stbDRs* are not equivalent and that the differences in the *stbDR* sequences of *stbS* have an impact on StbA binding, the ones carrying only two consensus sites and representing the wild-type configurations being more active.

Notably, additional band shifts migrating very slowly (above BI) are observed at the highest concentrations of StbA with stb2 (stb2, lanes 5 and 6). Such low mobility band shifts were also observed with substrates stb5, stb5a and stb5B. These might arise from interactions between two complexes CI, forming ‘sandwich complexes’, which would contain two DNA molecules. In this view, the two arrays of five *stbDRs* that compose *stbS* might be bridged together upon binding by StbA. To investigate such possibility, we employed the short–Long EMSA coupled to differential fluorescent DNA labeling method (26). We incubated StbA with a mixture of stb5a (CY5-labeled, 80 bp) and stb5b (CY3-labeled, 60bp) substrates. As shown in figure S5B (right panel), no additional complexes were detected in these conditions, thereby ruling out the possibility of StbA-bound ‘sandwich’ complexes composed of two distinct DNA molecules. The sandwich was neither detectable with stb5 mixed with stb1, stb2, stb3, nor with stb2’ substrates (Figure S5C).

Altogether these results demonstrate that StbA bound the *stbDR* region in a sequence-specific and concentration-dependent manner. They also strongly suggest that StbA binds cooperatively to 2 consecutive *stbDR* as a dimer and to 4 consecutive *stbDR* as two dimers, forming complexes with different stoichiometries.

### Specific DNA-binding activity of StbA to s*tbDR* resides in its N-terminal domain

To examine the role of the HTH DNA-binding motif contained in the N-terminal domain of StbA, EMSA were conducted using the StbA_1-75_ protein, a variant truncated for the C-terminal domain. As shown in Figure 5C, the StbA_1-75_ protein gave rise to similar patterns as those observed with StbA. No discrete band revealing stable complex formation was readily generated with a single *stbDR* (left panel, lanes 2-5). Interestingly, a slow-migrating complex was observed at high concentration of StbA (lane 6). As hypothesized above, this might correspond to interactions between complexes CI, although such ‘sandwiches’ could not be detected (figures S5B and C, and see above). Stable complexes equivalent to CI readily appeared with DNA substrates carrying 2 *stbDR* (stb2, central panel) and 3 *stbDR* (stb3, right panel). A second band corresponding to BI was also detected with stb3 (lanes 5 and 6). As observed with full-length StbA, incubation of StbA_1-75_ with stb5a generated mostly the complexes CII (Figure S5D, lanes 2-5). We thus concluded that the HTH motif included in the 75 N-terminal residues of StbA carries the specific and cooperative DNA binding activity of the protein.

### The transcriptional repressor activity of StbA is contained in its N-terminal domain

To investigate the role of the N-terminal domain of StbA in transcriptional repression, we performed transcriptional analyses of four StbA-repressed promoters containing 2, 3, 5 or 10 *stbDR* sequences (the *orf14*, *orf7*, *orf12,* and *stbA* of R388, respectively, Figure 1 and S6) in the presence of either StbA or StbA_1-75_. We compared the activity of these promoters in *E. coli* strains carrying a derivative of pUA66 vector (Zaslaver et al., 2006) that drives transcription of the *gfpmut2* reporter gene under the control of one of the *stbDR*-containing promoters, and a pBAD33 derivative expressing either StbA (pBAD33::*stbA*) or StbA_1-75_ (pBAD33::*stbA_1-75_*) (Fernandez-Lopez et al., 2014; Materials and Methods).

Repression rates were linearly related to the increase of the arabinose inducer and thus to the increase of StbA or StbA_1-75_ protein concentrations (Figure S6). As shown in Table 2, StbA_1-75_ repressed the activity of all four promoters, indicating that the StbA N-terminal domain contains the transcriptional repression activity of the protein.

**Table 2:**
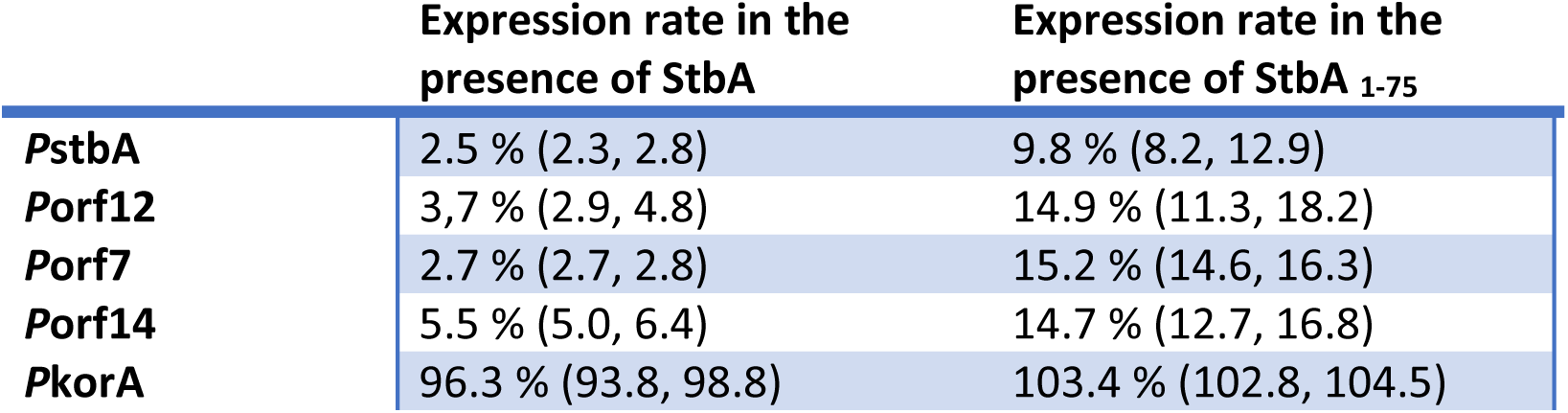
Transcriptional activity of stbDR-carriyng promoters in the presence of StbA or StbA _1-75_. Expression rates are expressed as the ratio between the expression levels (GFP/OD) measured in the presence and the absence of StbA or StbA_1-75_. StbA or StbA_1-75_ were produced from a co-residing plasmid pBAD33 (pBAD33::StbA1-75 or pBAD33::StbA1-75, respectively) and 2.5 10^-7^ of inducer (arabinose). Reference expression levels were measured in the presence of the empty vector (pBAD33). A control experiment with a promoter carrying no *stbDR* (pKorA) is shown. Extreme values are shown between brackets.

### Both domains of StbA are required for the control of partition and conjugative transfer of plasmid R388

To examine the role of the N-terminal domain of StbA on R388 stability and its interplay with StbB activity *in vivo*, we constructed a variant of R388 carrying a truncated *stbA* gene that encodes StbA_1-75_ (R388*stb*A_1-75_, Materials and methods). We first measured plasmid loss rates from serial cultures in non-selective medium, and some of the results are shown in Figure 6A. R388 wt was highly stable in these conditions, while R388Δ(*stb*A) was lost at an average rate of 2.5 ± 0.7 % per cell per generation (Figure 6A). R388*stb*A_1-75_ was also lost, although at a slightly lower rate (with an average loss of 1.3 ± 0.6 %). This result indicated that StbA N-terminal domain stabilizes R388 only partially *vivo*.

**Figure 6.**
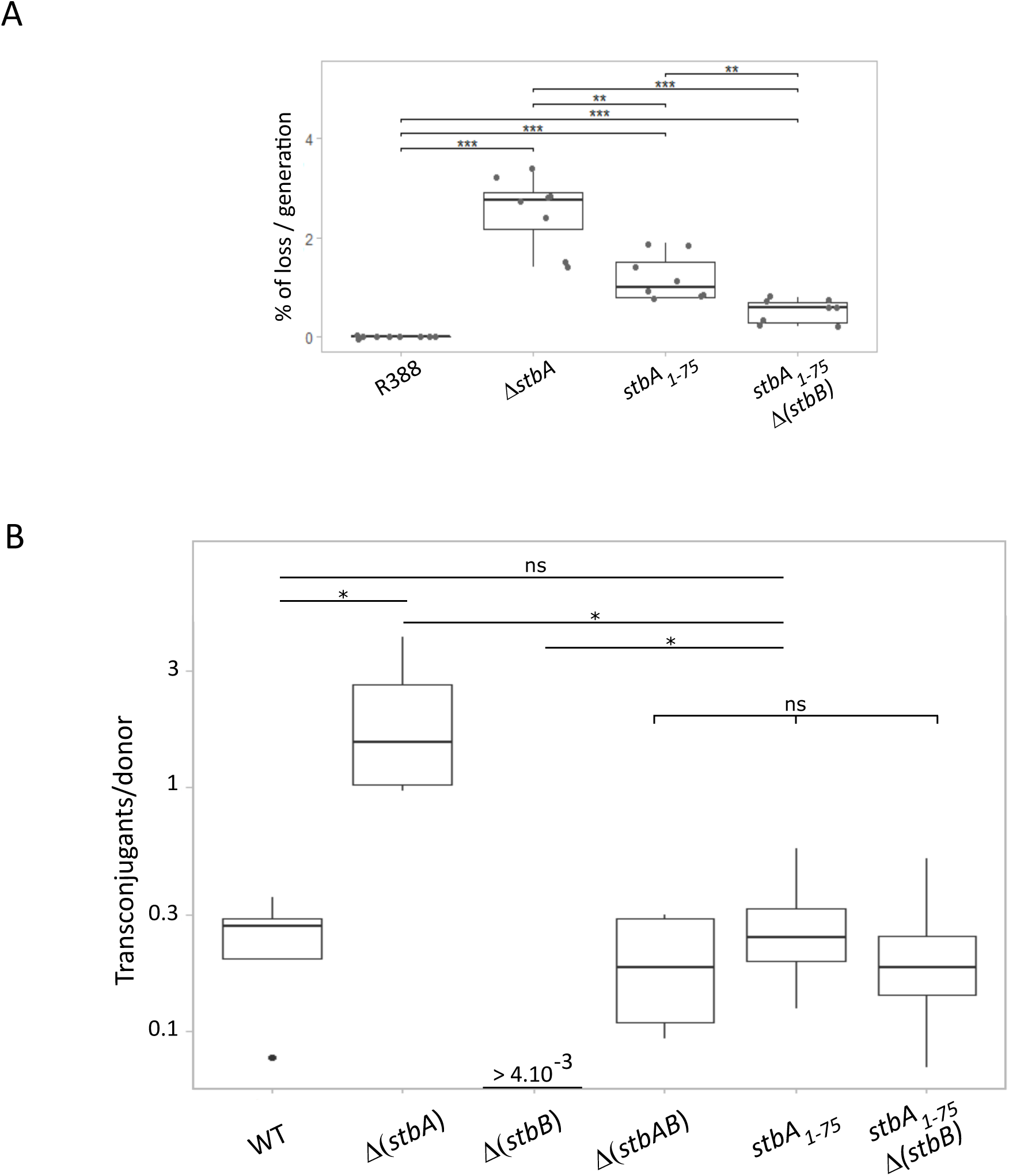
The DNA-binding domain of StbA shows partial activity on the stability and in the control of conjugation of plasmid R388. The stability (**A**) and the conjugation frequencies (**B**) of different derivatives of plasmid R388 are shown. Plasmids are indicated on the X-axis. **A.** Stability was measured as the rate of loss per cell per generation from strain LN2666 (Materials and Methods). **B.** Conjugation frequencies as the number of transconjugants per donor (a Log scale is used for the Y-axis). Error bars show standard deviations calculated from at least four independent assays.

Thus, while the StbA N-terminal alone can stabilize R388 to a certain extent, its C-terminal domain is required for full stability. We then repeated the same experiments in absence of StbB, which is not involved in R388 wt stability (5). As previously shown, R388Δ(*stbB*) was as stable as R388 wt, while R388Δ(*stbAB*) was as unstable as R388Δ(*stbA*) ((5); R388Δ(*stbAB*), 2.3 ± 0.5 %, not shown in Figure 6A). Surprisingly, the instability provoked by the *stbA_1-75_* mutation was partly suppressed by *stbB* deletion (R388*stb*A_1-75_ Δ(*stbB*), 0.5 ± 0.2 %). This may be linked to a weaker nucleoid-associated positioning of R388*stbA_1-75_* compared to R388 wt that would render the former more sensitive to a nucleoid exclusion activity of StbB (see below).

StbA was previously found to inhibit or even to fully abolish conjugative transfer in the presence or absence of StbB, respectively (5). In contrast, StbB has been shown to stimulate conjugation whether StbA is present or not, whereas the absence of both StbA and StbB had no effect on conjugation frequencies (5). As confirmed in Figure 6B, while R388Δ(*stbAB*) exhibited similar conjugation frequencies as R388 wt, inactivation of StbA led to a significant increase of conjugation frequencies compared to R388 wt, whereas no conjugation was detected when StbB was inactivated (frequency < 4.10^-3^). The mechanism by which StbA inhibits conjugation is only partially understood. It correlates with StbA-dependent positioning of plasmid copies exclusively in the nucleoid area, which is detrimental for conjugation (5). To study the role of the N-terminal domain of StbA in the control of conjugation, we measured the conjugation frequencies of R388*stbA_1-75_*. Interestingly, R388*stbA_1-75_* conjugation frequency was similar to that of wild-type R388, but significantly lower than that of R388Δ(*stbA*) (Figure 6B). This suggested that StbA_1-75_ retained the activity limiting conjugation efficiency. Yet, contrary to wt, inactivation of StbB in the R388*stbA_1-75_* did not affect its conjugation frequency (R388stbA_1-75_Δ(*stbB*), Figure 6B). Thus, StbA_1-75_ has lost the ability to inhibit conjugation when StbB is absent.

### The StbA C-terminal domain is required for subcellular localization of plasmid R388

To determine whether R388*stbA_1-75_* instability is correlated with an abnormal subcellular positioning, as previously shown for R388Δ(*stb*A), we analyzed the subcellular localization of the plasmid in live *E. coli* cells using the *parS*/ParB-GFP system (5). Figure 7A shows representative images of the location of GFP foci position as a function of the cell length. R388*stbA_1-75_* foci showed a tendency to localize at the cell center and polar regions, with a pattern resembling that of R388Δ(*stb*A). To compare the localization profile of R388*stbA_1-75_* with R388 and R388Δ(*stb*A), we analyzed the distribution of GFP foci in the length of half-cells divided into five equal sections from one pole to mid-cell (Figure 7B). R388*stbA_1-75_* foci were found mainly at the cell center (33% of foci at 0.4 to 0.5 fractional cell length compared to 27% for R388) and within the most polar region (24% of foci at 0 to 0.1 fractional cell length compared to 12.5% for R388). However, χ^2^ test revealed that these differences were not highly significant (χ^2^=7.96, p-value=0.092). Interestingly, R388Δ(*stb*A) and R388*stbA_1-75_* distributions were also not significantly different (χ^2^=4.2, p-value=0.380), while R388Δ(*stb*A) and R388 were (χ^2^=10.9, p-value=0.027). These data indicated that R388*stb*A_1-75_ displays partial activity in the cellular positioning of plasmid copies, possibly correlated with plasmid instability but less severe than that of R388Δ(*stbA*). Noteworthy, R388*stbA_1-75_* foci showed a tendency to localize less frequently at the most polar region of the cell than R388Δ(*stb*A) foci (42% compared to 57%, respectively, at 0 to 0.2 fractional cell length), but more at mid-cell (23% compared to 33%, respectively, at 0 to 0.2 fractional cell length). In contrast to R388Δ(*stbB*) foci, which were previously shown to localize exclusively to nucleoid areas and not at the cell poles (5), the localization pattern of R388Δ*stbA_1-75_*(*stbB*) was similar to that of R388Δ*stbA_1-75_*. Thus, plasmid localization correlated with conjugation efficiency, as deletion of *stbB*, which leads both to retention of plasmid copies in the nucleoid and abolition of transfer in the presence of StbA (5), did not affect conjugation frequencies in the R388Δ*stbA_1-75_* mutant compared to R388 wt (Figure 6B).

**Figure 7.**
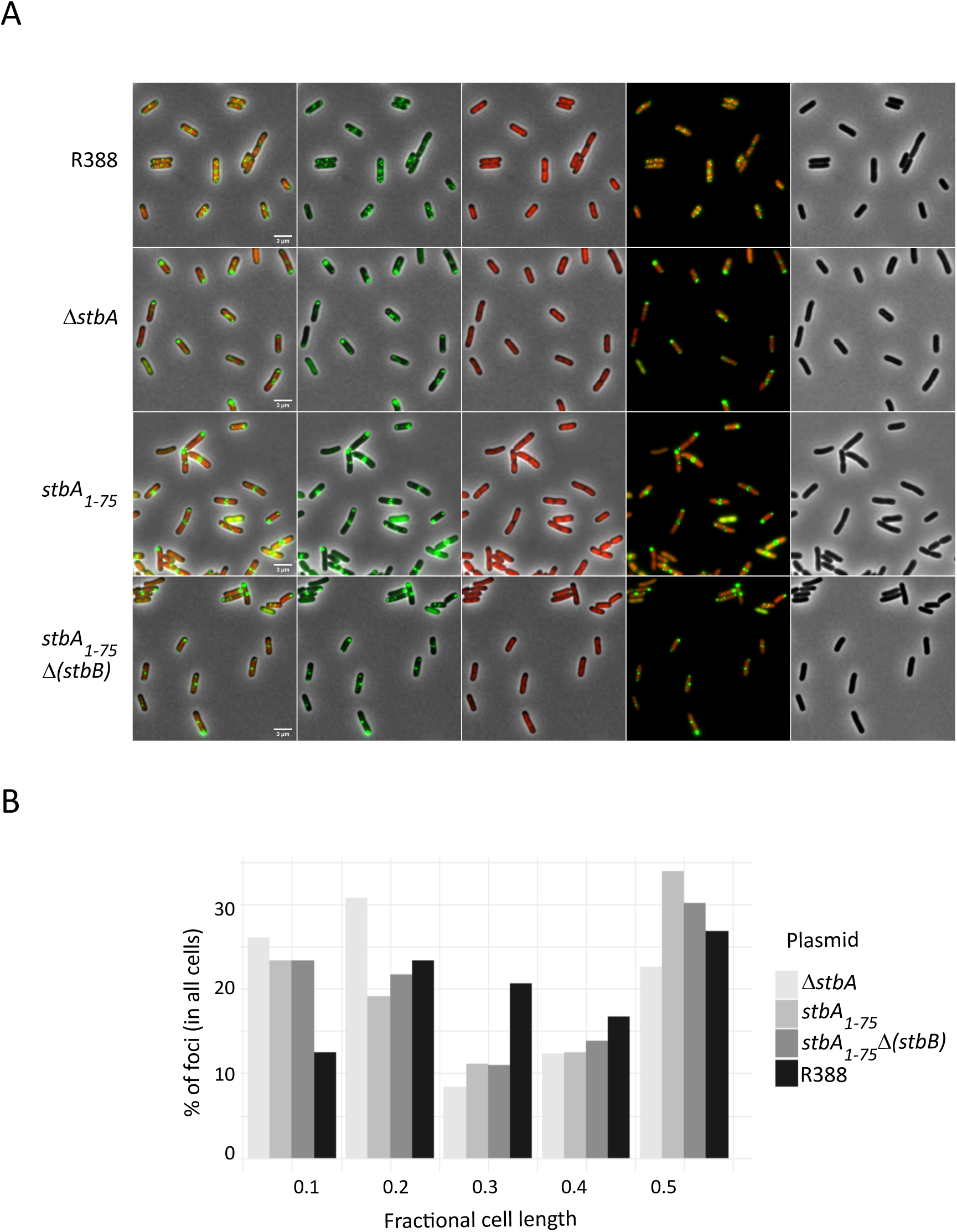
The DNA-binding domain of StbA shows partial activity on subcellular localization of plasmid R388. **A.** Live cell imaging of *E. coli* strain LN2666 containing R388 derivatives harboring *parS* (R388, R388Δ*stbA*, R388*stbA_1-75_* and R388*stbA_1-75_*Δ*stbB*) and expressing GFP-D30ParB from plasmid pALA2705. Scalebar = 3 μm. From left to right, panels show fluorescence pictures of (i) ParB-GFP-tagged plasmids (green) merged with fluorescence pictures of HU-mcherry (red) and the phase contrast pictures, (ii) ParB-GFP alone, (iii) HU-mcherry alone, (iv) ParB-GFP-tagged plasmids merged with HU-mcherry, and (v) phase contrast alone. **B.** Distribution of GFP foci within the different fractions of cell length. The distance of foci to the closest cell pole was measured and sampled into five equal sections of cell length from the pole to mid-cell (R388 n= 5016, R388Δ*stbA* n= 2686, R388*stbA_1-75_* n= 2405, R388*stbA_1-75_*Δ*stbB* n= 2383).

Altogether, our data suggest that, while StbA N-terminal DNA-binding domain displays partial activity, both domains of StbA are required for proper positioning of R388 copies, correlating with plasmid stability and the control of conjugation.

## DISCUSSION

The segregation system of plasmid R388 differs from the bacterial systems described, in that it consists only in a single centromere-binding protein, StbA, to achieve effective partition of a low-copy-number plasmid. StbA is multifunctional, since it also plays a key role in the regulation of the expression of several R388 genes and in the control of conjugation. Our study provides the first structural and biochemical report of a member of the StbA family of proteins. We show that StbA is a two-domain protein, of which the N-terminal domain, StbA_1-75_, contains an HTH motif that supports all the specific DNA-binding activity of the protein, and correlate this with StbA activities *in vivo*.

StbA activities involved in the stable inheritance and in the inhibition of conjugation of plasmid R388 have previously been correlated with the confinement of plasmids at nucleoid areas (5). Our data indicate that its N-terminal domain displays partial activity that also correlate with the subcellular positioning of the plasmid. First, we show that R388*stbA_1-75_* is unstable but to a lesser extent than R388Δ(*stbA*) (Figure 6A), which correlates with the sub-cellular positioning of R388*stbA_1-75_* being in-between that of the plasmids R388 and R388Δ(*stbA*) (Figures 6B and 7). Deletion of the C-terminal domain of StbA could thus alter the interactions between StbA and the nucleoid, resulting in poor retention of plasmids at the nucleoid and therefore increased plasmid loss. Second, our results indicate that R388*stbA_1-75_* conjugation frequencies are similar to that of R388 wt, and much lower to that of R388Δ(*stbA*), demonstrating that StbA_1-75_ retains the ability to inhibit conjugation (figure 6). However, StbB, which is strictly required for conjugation in the presence of StbA, is not essential in the absence of the StbA C-terminal domain. Again, this correlates with plasmid localization since conjugation events can be attributed to the copies of the plasmid not being effectively maintained at the nucleoid by StbA_1-75_. However, there are twice as many copies of R388*stbA_1-75_* as R388 at the cell poles (figure 7), and one could expect higher levels of transfer of R388*stbA_1-75_*, as observed for R388Δ(*stbA*). This indicates that the control of conjugation can probably not only be attributed to the subcellular positioning of the plasmids, but also to the binding of StbA to R388 *per se*. In this view, StbB could interact with C-terminal domain of StbA to stimulate conjugation by releasing the plasmid from StbA. We conclude that, whatever the way StbA positions plasmid copies at the nucleoid, its C-terminal domain is likely to be required and could interact with StbB to control conjugation. Third, we demonstrate that StbA_1-75_ exhibits partial repression activity on the *stbDR*-bearing promoters of several R388 genes, suggesting that the C-terminus of StbA might stabilise interactions between StbA and the *stbDR* sites (Table 2 and Figure S6).

The StbA N-terminal domain includes a well-conserved DNA-binding domain that folds into an HTH motif not predicted *in silico* based on sequence homology and structure prediction programs. StbA HTH motif consists of the widely spread basic tri-helical version of the HTH domain, found in many proteins, such as bacterial transcription factors of the FIS family, or the eukaryotic Homeo, POU and Myb domains (29). StbA N-terminal domain is structurally most related to the HTH motif of the protein Rv3488 (31), which belongs to the PadR subfamily II of transcription regulators, including small proteins, like StbA, of ∼ 110 amino acids protein, such as LmrR (36), BcPadR1 and BcPadR2 (32). These proteins harbor a conserved HTH motif, called winged-HTH because it comprises at the C-terminus of the recognition helix α3, two β-strands devoted to bind with the operator DNA, and a short C-terminal domain containing a single α-helix (Figure 4B). Like most bacterial transcription factors, structurally characterized PadR subfamily II proteins form dimers via a two-fold symmetry in the crystal. In the dimeric structures, the C-terminal helix flanking the winged HTH motif of one monomer interacts with structural parts of the HTH motif of the other monomer ((31); (32); (43) ; (36) ; (44) ; (45)).

Our data *in vitro* indicate that StbA binds specifically to the *stbDR* sequences with a strong cooperativity that lead to the binding of one StbA dimer to every two *stbDR* (Figures 5 and S5). Full-length StbA is indeed dimeric in solution, but, in contrast, StbA N-terminal domain (StbA_1-75_) is monomeric (Figure S2). Yet, StbA_1-75_ shows similar specific and cooperative binding patterns to the *stbDR* as the full-length StbA, suggesting that interactions between StbA_1-75_ monomers may be promoted by binding to DNA. Although binding cooperativity to DNA often involves direct protein–protein interactions, binding of StbA_1-75_ monomers could be promoted by local protein-induced changes in DNA shape that would create an optimized DNA conformation for binding of the second protein (46). Also, our structural data reveal a putative dimerization interface involving hydrophobic interactions between α1 of one monomer and α2 of the other monomer (Figure S3), which is consistent with our bacterial two-hybrid assays results showing that StbA N-terminal domain interacts with itself *in vivo* (Figure S2). Altogether, these observations suggest that StbA C-terminal part, unlike PadR proteins, may not be involved in oligomerization, or could just have a role in stabilizing StbA multimers.

By analogy to all plasmids and chromosomal CBPs described so far, StbA might assemble into high-order complexes at its centromere-like site. Our results indicate that *in vitro*, StbA and StbA_1-75_ form specific high molecular weight complexes. These could arise from the association of monomers of the StbA N-terminal domain to form high-order multimer through intermolecular hydrophobic interactions and hydrogen bonds, as suggested by the structural data. Another possibility could be the assembly of complexes comprising different pieces of *stbDR*-carrying DNA, as might be evoked by the organization of *stbS* in two arrays of 5 *stbDR*. Our EMSA did not allow to detect complexes containing two separate DNA molecules carrying *StbDR* sites. These data however do not rule out the possibility the formation of a sandwich complex that bring together the two *stbDR* arrays of *stbS* inside the same DNA molecule *in vivo*.

*StbDR* sequences are spaced with 2 bp, which forms an 11-bp cycle corresponding to a full helix turn, suggesting that the StbA dimers bind to the same side of the DNA. This has been observed for the CBP ParR of plasmids pB171 and pSK41, which assemble in a cooperative manner in a continuous structure on their cognate centromere that is organized very similarly to *stbS* ((47); (48)). *StbS* organization and its location upsteam the *stb* operon, as well as the fact that StbA carries an HTH DNA-binding domain are also reminiscent characteristics of type III partitioning systems, of which the most studied system is *tubRZC* of *Bacillus thuriengensis* plasmid pBtoxis ((49);(2)). According to this, StbA may form a large DNA-protein filament structure around the two sets of *stbDR* of *stbS*, as observed for TubR proteins around the seven 12-bp direct repeats arranged in two sets forming *tubC* ((50) ; (51); (52)). Similarly, the CBP AspA of the archeal plasmid pNOB8, which shows like StbA homology with the PadR family of transcriptional regulators, spreads onto DNA forms a protein-DNA superhelical structure (38).

As all the plasmid *par* operons, the *stb* operon is autoregulated. StbA regulates the expression of its own promoter and also four others promoters of genes located in the maintenance region of plasmid R388 (7). We show here that StbA forms stable specific complexes with DNA substrates carrying at least two direct repeats of *stbDR in vitro*. This might be biologically relevant since StbA operator regions consist in arrays of two, three or five direct repeats of *stbDR*, with the exception of *stbS*, which contains ten *stbDR* arranged in two sets in the *stbA* promoter. This dual role of StbA to act as a transcriptional repressor of its operon, and of other unrelated genes, and also as R388 CBP, raises the question of how *stbS*, which is strictly required for plasmid stability, is recognized by StbA as a segregation site. The CBP of plasmid RK2, KorB, also regulates many genes on the plasmid, including the partition genes *incC* and *korB*. KorB recognizes and binds to a palindromic operator found 12 times on plasmid RK2 (OB1-OB12). In contrast to *stbS*, the OB3 site, is required for RK2 partition but is not involved in the regulation of the expression of the partition genes, which is achieved through KorB binding to OB1, flanking the −35 sequence (53). Previous study showed that the presence of the other *stbDR* in the promoter of several R388 genes is not sufficient to ensure R388 stability, indicating that this arrangement in two sets of *stbDR* arrays is important for segregation (5). Notably, the loss rates of R388 deleted of *stbS* is similar to that of plasmid R388*stbA_1-75_*, which might suggest that the formation of the segregation complex requires the arrangement of the two sets of *stbDR* of the *stbS* and requires the C-terminal domain of StbA.

We aim to understand the trade-offs resulting from the multiple roles of StbA in plasmid R388 physiology. In contrast to typical partition operons, the other genes of the *stb* operon, *stbB* and *stbC*, are not required for plasmid stability. We speculate that the ATPase StbB could counteract StbA activity by releasing R388 copies from the nucleoid through interactions with StbA and/or other factors. Conjugation and stability seem to be competing mechanisms in plasmid R388 molecular physiology. We speculate that the R388 segregation system involving a single centromere-binding protein might represent an ancestral mechanism, from which the typical Par systems originated. These might have co-opted a dedicated NTPase to improve partition by moving the partition complexes polewards and thus make partition and conjugation more uncoupled processes.

Putative host partners of the StbA/*stbS* system and the associated mechanism for R388 segregation are currently unknown, as well as the way by which StbB controls conjugation. This knowledge will be crucial to understand how StbA controls vertical and horizontal transfer of plasmid R388. PTU-W plasmids, including the three typical members R388, pSa and R7K are among the broadest host range plasmids in Proteobacteria. Their successful transfer and stable inheritance have been reported in many bacterial species, including all the ‘ESKAPE’ pathogens ((54); (8)). These plasmids are good examples for the acquisition of antibiotic resistance genes as a consequence of the pressure exerted by antibiotic usage (55). Our studies point out that a single mutation in the *stbA* gene could generate plasmids with much higher transfer ability (5), thus alarming on the possible emergence of such ‘superspreader’ plasmids. It has recently been reported that an effective approach to limit the spread of antibiotic resistance genes would be the combination of the control of plasmid transmission by conjugation (reviewed in (56) and (57)) and fostering plasmid loss (58). In this view, the StbAB system, which controls the interplay between plasmid conjugation and segregation may be an interesting target. Understanding the mechanisms of integration of vertical and horizontal modes of plasmid propagation within bacterial populations is of utmost importance given that they are major contributors to the spread of antibiotic resistance (59).

## Data Availability

Atomic coordinates and structure factors for the reported crystal structure of StbA have been deposited in the Protein Data bank with accession number 7PC1.

## Supplementary data

### Supplementary materials and methods

Bacterial-two-hybrid analysis (BACTH).

Size exclusion chromatography

## Fundings

This work was supported by the Spanish Ministry of Economy, Industry and Competitiveness grant BFU2017-86378-P to F.dlC., by the Spanish Ministry of Science (MCI/AEI/FEDER,UE) grant PGC2018-093885-B-I00 to G.M., by French National Research Agency, grant numberANR-18-CE35-0008 to J.-Y.B. and by University P. Sabatier grant to C.G..

## Acknowledgments

Structural experiments were performed at PROXIMA beamline at the SOLEIL Synchrotron (France) with the collaboration of Beatrix Guimaraes. We thank all members of the GeDy team for fruitful discussions.

## Supplementary data

### Supplementary materials and methods

Bacterial-two-hybrid analysis (BACTH). Dimerization of StbA and StbA_1-75_ was analyzed *in vivo* using the bacterial two-hybrid system ((59); (60)) in the *E. coli* BTH101 *cyaA* strain. N- and C-terminal CyaAT18 or CyaAT25 fusions of *stbA* or *stbA_1-75_* were constructed using plasmid pKT25, pUT18C and pUT18. *stbA* and *stbA_1-75_* genes were amplified by PCR from plasmid R388 using primers G158 and G159, and G158 and G254, respectively and cloned between the *BamH*I and *EcoR*I restriction sites of pUT18C and pKT25, or using primers G161 and G162, and G161 and G256, respectively and cloned between the *BamH*I and *Hind*III restriction sites of pUT18.

Derivative BACTH vectors were co-transformed into *E. coli* BTH101 *cyaA* in all pairwise combinations. Several colonies of co-transformants were selected and grown in LB medium supplemented with 100 μg/ml ampicillin, 50 μg/ml kanamycin and 0.5 mM IPTG overnight at 30°C. Overnight cultures were then spotted on MacConkey plates with maltose as a carbon source containing 100 μg/ml ampicillin, 50 μg/ml kanamycin and 0.5 mM IPTG and plates were incubated at 30°C during 48 h.

### Size exclusion chromatography

Analytical gel filtration experiments were performed at 4°C using a Superdex 75 (10/300 GL) (GE Healthcare) with the FPLC system (Amersham Biosciences). Samples (250 µl) were injected onto the column pre-equilibrated with 20 mM Tris pH 7.5, 200 mM NaCl, 2 mM EDTA and 1 mM DTT and run over with the same buffer at a rate of 0.5ml/min and monitored by absorbance at 280 nm. For size estimation, the column was calibrated with ovalbumin (42.7 kDa), ribonuclease A (13.7 kDa) and Aprotinin (6.5 kDa). The K_av_ value was calculated for each standard protein (using the equation (V_e_ - V_0_)/(V_c_ - V_0_), where V_e_ is the elution volume for the protein, V_0_ the column void volume (V_0_ = 8 ml) and V_c_ the geometric column volume (V_c_ = 24 ml)), and plotted against the logarithm of standard molecular weights.

## Supplementary figures

**Table S1.** Strains and plasmids.

| Strains | Genotype/relevant properties | AB- res | Source/reference |
| --- | --- | --- | --- |
| DH5α | F<sup>-</sup> *endA1 recA1 gyrA96 thi-1 hsdR17 supE44 relA1 Δ(argF- lacZYA) U169 Φ80d lacZ DM15 gyrA96* | Nx | Sambrook et al., 1989 |
| LN2666 | F<sup>-</sup> W1485 *thiA thyA leu deoB rpsL* | Sm | Cornet et al., 1994 |
| LN2666 HU-mcherry | LN2666 containing a *hupA::mcherry* translational fusion that expresses the nucleoid associated protein HU fused with mCherry to label chromosomal DNA. | Sm | This study |
| BW27783 | BW25113 DE(*araFGH*) F(*ΔaraEp* P<sub>CP8</sub>-*araE*) | Rif | Khlebnikov et al., 2001 |
| DY380 | DH10B derivative containing a defective λ prophage; *red*, *bet* and *gam* genes are controlled by the temperature-sensitive λcl857 repressor | Sm | Lee et al., 2001 |
| C41 (DE3) | F<sup>-</sup> *ompT dcm hsdS* (r<sub>B</sub>- m<sub>B</sub>-) *galλ* (DE3) |  | Miroux and Walker, 1996 |
| **Plasmids** |  |  |  |
| R388 | R388 *parS-Cm* | Tmp,Cm | Guynet et al., 2011 |
| R388Δ*stbA* | R388 *parS-Cm Δ(stbA)* | Tmp,Cm | Guynet et al., 2011 |
| R388Δ*stbB* | R388 *parS-Cm Δ(stbB)* | Tmp,Cm | Guynet et al., 2011 |
| R388Δ*stbABC* | R388 *parS-Cm Δ(stbABC)* | Tmp,Cm | Guynet et al., 2011 |
| R388*stbA*<sub>1-75</sub> | R388 *parS-Cm* carrying a truncated *stbA* gene (*stbA*<sub>1-75</sub>) that encodes StbA<sub>1-75</sub> | Tmp Cm Km | This study |
| R388 *stbA*<sub>1-75</sub>Δ*stbB* | R388Δ*stbB* carrying a truncated *stbA* gene gene (*stbA*<sub>1-75</sub>) that encodes StbA<sub>1-75</sub> | Tmp Cm Km | This study |
| pAPT110 | Used for p4G39 and p4G41 constructions (Rep p15A) | Km, Sp/Sm | Polard and Chandler, 1995 |
| p4G39 | pAPT110 carrying the *stb* operon with the truncated *stbA*<sub>1-75</sub> | Km, Sp/Sm | This study |
| p4G41 | pAPT110 carrying the *stb* operon with the truncated *stbA*<sub>1-75</sub> and deleted of *stbB* | Km, Sp/Sm | This study |
| pET-StbA | pET29C (Novagen) derivative producing StbA | Km | This study |
| pET-StbA<sub>1-75</sub> | pET29C (Novagen) derivative producing StbA | Km | This study |
| pBAD33 | Used for pBAD-StbA and pBAD-StbA<sub>1-75</sub> constructions (Rep p15a) | Cm | Guzman et al., 1995 |
| pBAD-StbA | pBAD33 derivative carrying *stbA* between *XbaI* and *HindIII* sites | Cm | Fernandez-Lopez et al., 2014 |
| pBAD-StbA<sub>1-75</sub> | pBAD33 derivative carrying *stbA*<sub>1-75</sub> between *XbaI* and *HindIII* sites | Cm | This study |
| pALA2705 | *GFP-ΔN30parB* expressing from *Plac* promoter | Ap | Li and Austin, 2002 |

**Table S2.**
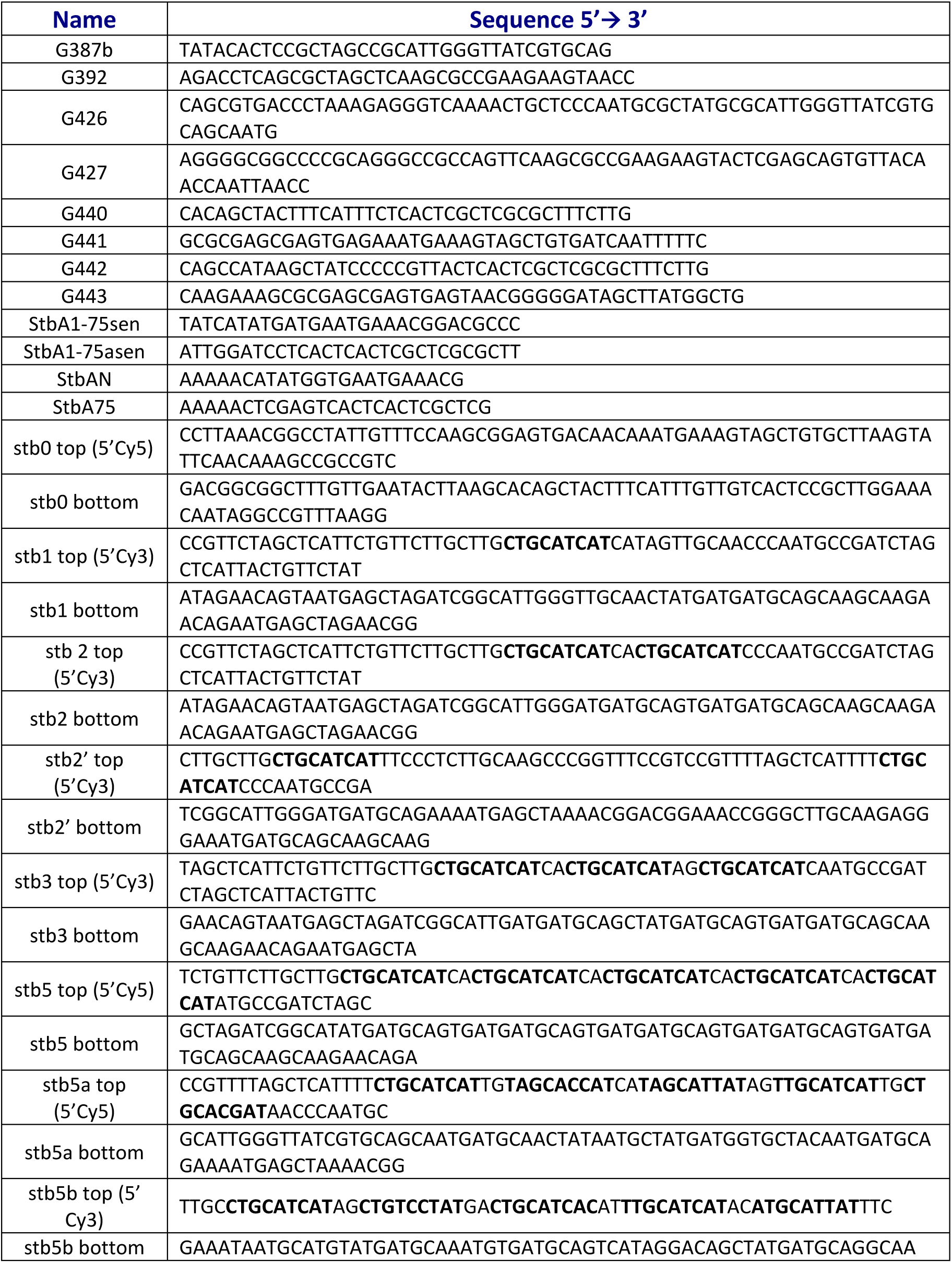
Oligonucleotides used in this study.

**Table S3.**
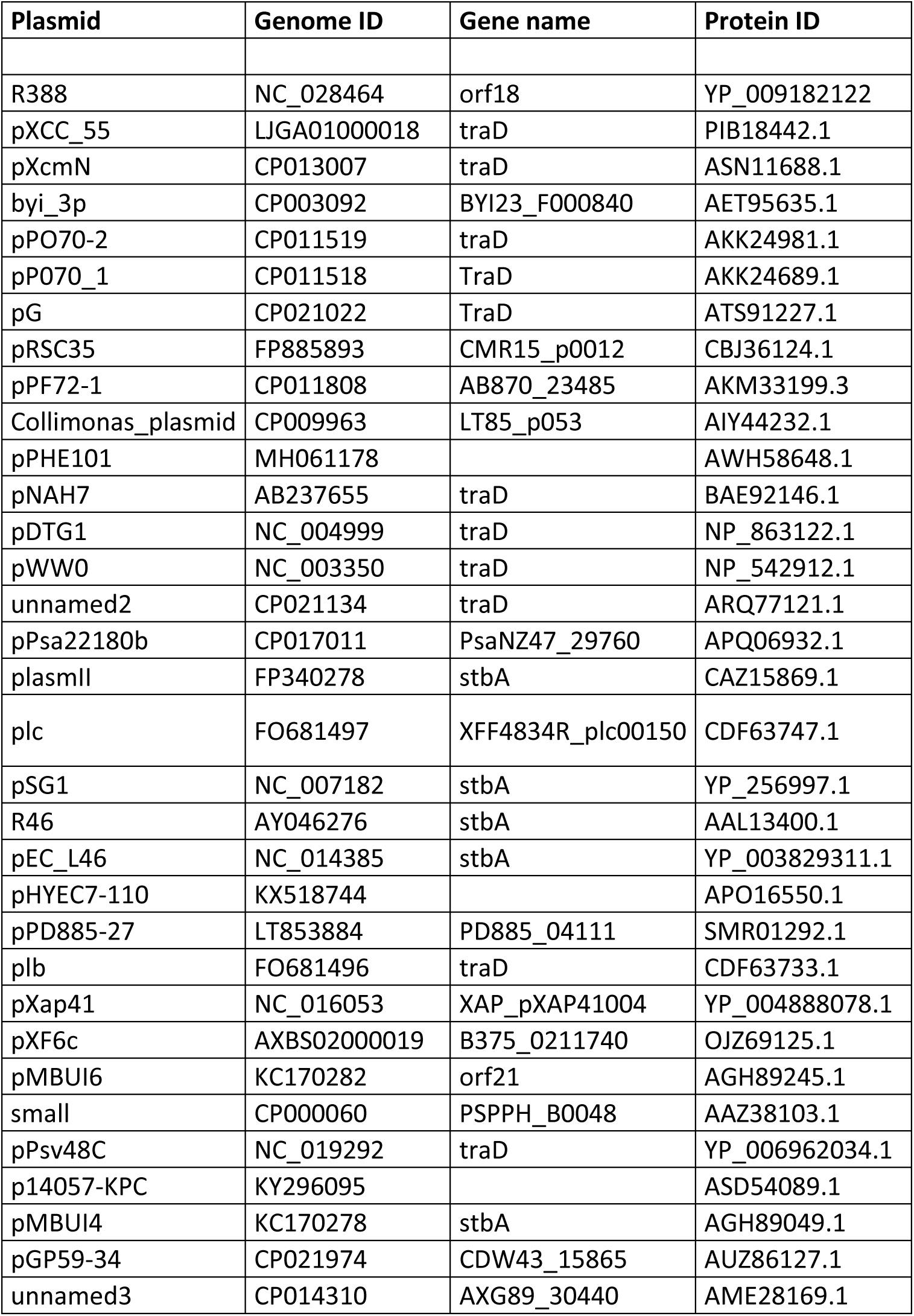
Sequence ID of proteins used in Figure 2.

**Figure S1.**
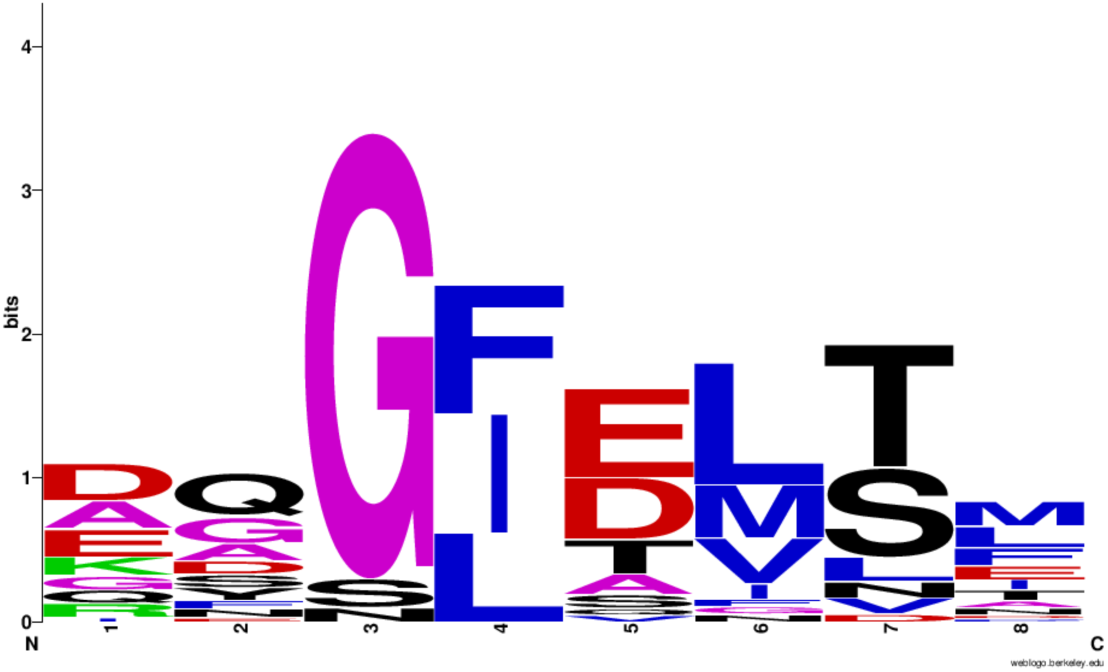
WebLogo of amino-acid sequences of the turn between α2 and α3 of StbA proteins. The sequence logos of the turn between α1 and α2 helices of StbA proteins was generated using the WebLogo software (61) from amino-acid sequences of StbA proteins shown in Figure 2. Hydrophobic, acidic, small, basic and polar non-charged residues are shown in blue, red, purple, green and black, respectively.

**Figure S2.**
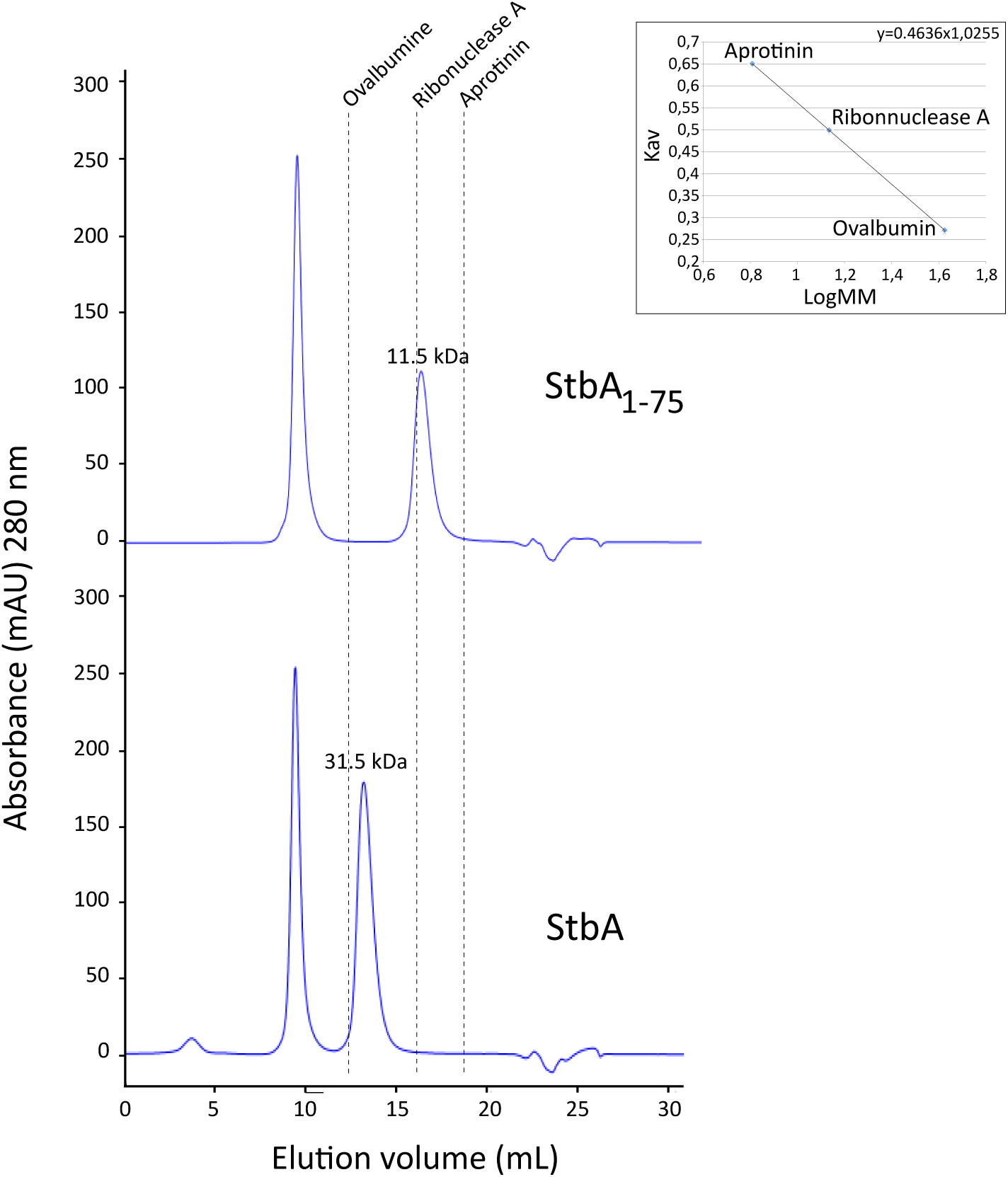
Gel filtration chromatography analysis of purified StbA and StbA_1-75_. StbA and StbA_1-75_ were analyzed at 60 μM. Elution profiles were monitored by absorbance at 280 nm. The elution positions of molecular weight standards (Ovalbumin (43 kDa), Ribonuclease A (13.7 kDa) and Aprotinin (6.5 kDa) are indicated. The molecular mass expected for a monomer is 17 kDa for StbA and 9.9 kDa for StbA_1-75_. StbA and StbA_1-75_ eluted at 16.5 ml and 13.25 ml, respectively. The experimental Kav values suggest a molecular mass in solution of 31.5 kDa for StbA and 11.5 kDa for StbA_1-75_ corresponding to a dimer and to a monomer, respectively. In both cases, the pick corresponding to high molecular weight species (void fractions) was likely due to protein aggregation during storage.

**Figure S3.**
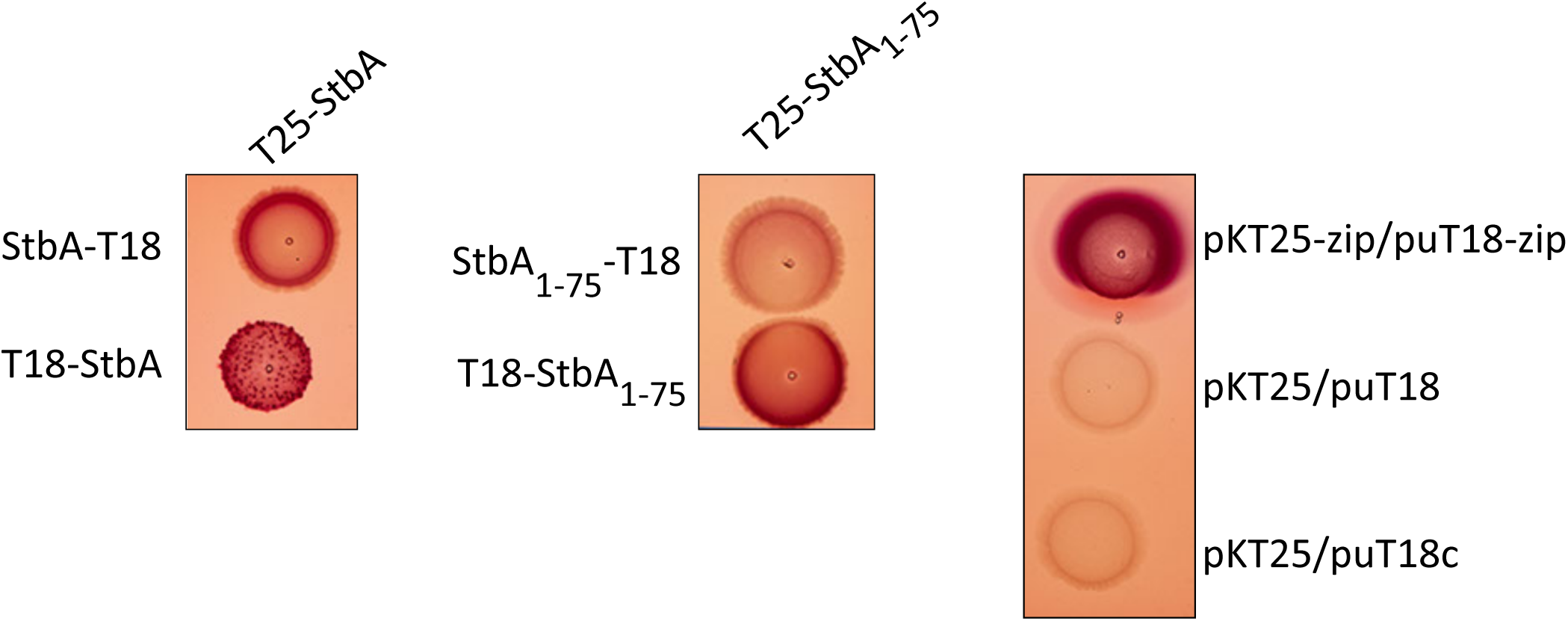
StbA and StbA_1-75_ dimerization *in vivo* in the BACTH system. Bacterial two-hybrid of StbA and StbA_1-75_ tagged at their N-terminus (T25-StbA, T18-StbA and T25-StbA_1-75_, T18-StbA_1-75_) or C-terminus (StbA-T18 and StbA_1-75_-T18). Double transformants of *E. coli* BTH101 *cyaA* with compatible plasmids encoding CyaA fragment T18 or T25 fused to either StbA or StbA_1-75_ were analyzed on indicator MacConkey plates with maltose as a carbon source. Purple spots are indicative of interactions between interactive proteins. Image representative of 3 independent trials.

**Figure S4.**
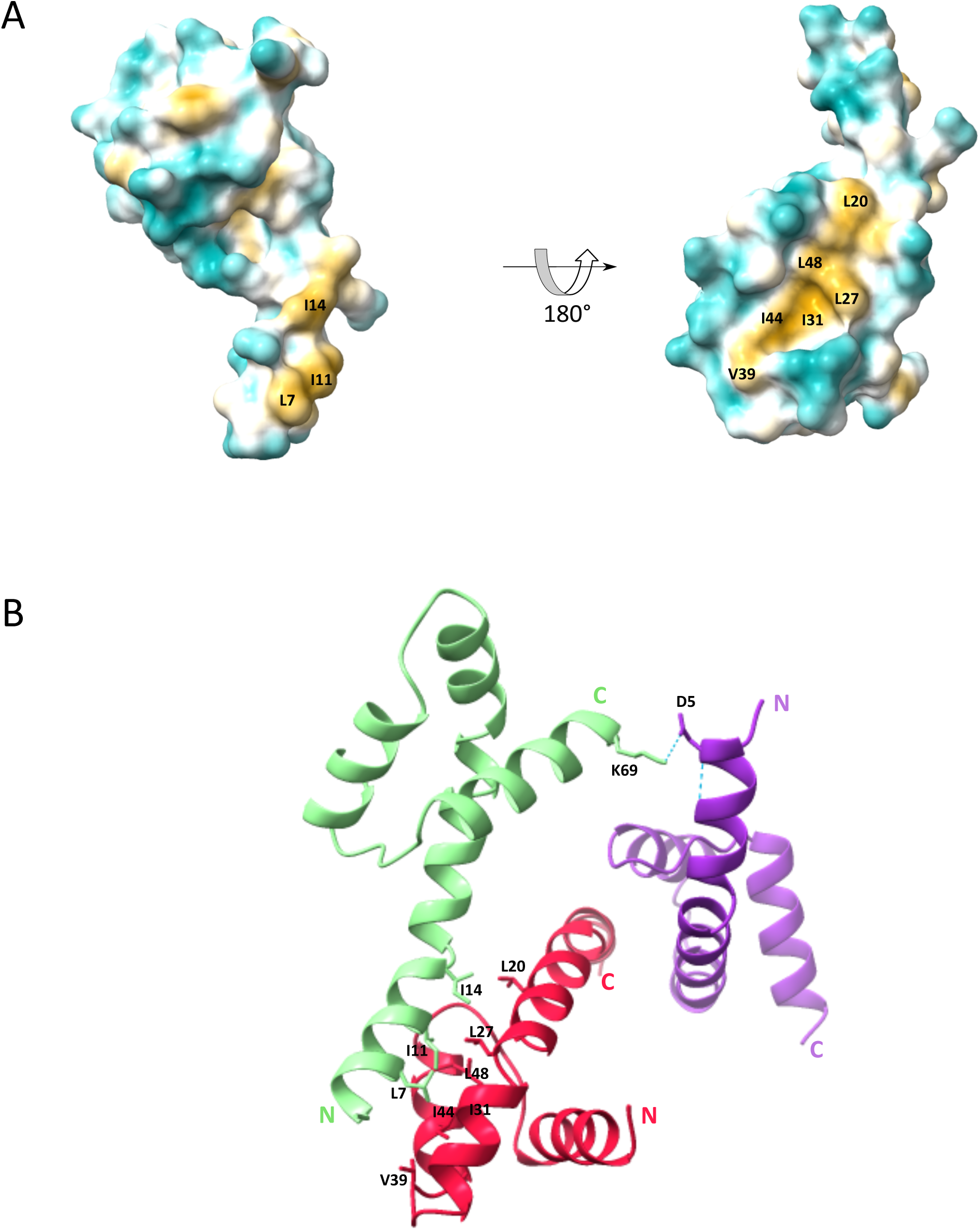
Crystallographic analysis of StbA multimerization. A. The molecular surface map of StbA_1-75_ reveals a patch of hydrophobic residues, of which several are conserved or with similar side chains, as seen in the sequence alignment (Figure 2), and which are putatively involved in dimerization. Coloring is from dark cyan for most hydrophilic through white to dark goldenrod for most hydrophobic. B. Ribbon diagram of a trimer of StbA_1-75_ as observed in the crystal showing a three-fold symmetry between StbA_1-75_ subunits. The hydrogen bond (dotted blue line) formed by Asp_5_ from monomer 1 (green) and Lys_69_ from monomer 3 (purple) and that may stabilize the trimer is indicated.

**Figure S5.**
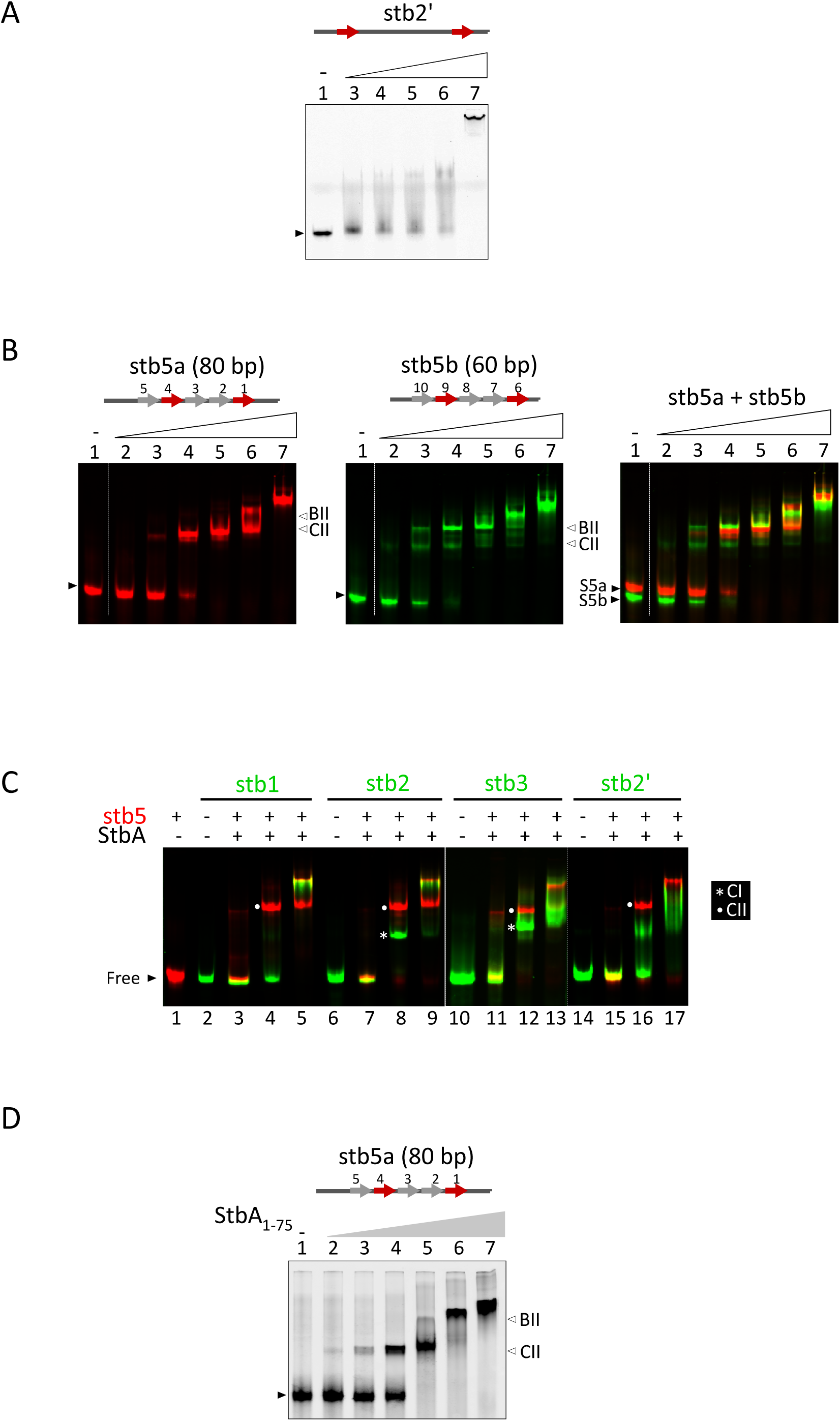
DNA-binding activities of StbA and StbA_1-75_ by electrophoretic mobility shift assay (EMSA). EMSA performed on 5% polyacrylamide gels. DNA substrates are schematized at the top of the figure. Free DNA and high molecular weight complexes are indicated by black wedges and a vertical bar, respectively. A. A Cy3-labeled 80-pb DNA fragment carrying 2 *stbDR* consensus sequences separated by 43 bp (stb2’) was incubated with 0 (lane 1), 1 μM (lane 3), 2 μM (lane 4), 4 μM (lane 5), 8 μM (lane 6), and 16 μM (lane 7) StbA. B. stb5a (Cy5-labeled, left panel), stb5b (Cy3-labeled, central panel) or both (right panel) substrates were incubated with 1 μM (lanes 3, 7, 11, 15), 4 μM (lanes 4, 8, 12, 16), 16 μM (lanes 5, 9, 13, 17) StbA. C. stb5 substrate (Cy5-labeled) was incubated with 1 μM (lanes 3, 7, 11, 15), 4 μM (lanes 4, 8, 12, 16), 16 μM (lanes 5, 9, 13, 17) StbA in the presence of a second Cy3-labeled DNA fragment carrying 1 (stb1), 2 (stb2 and stb2’) or 3 *stbDR* (stb3). CI and CII complexed are indicated by white stars and circles, respectively. D. stb5a (Cy5-labeled) was incubated with 0 (lane 1), 500 nM (lane 2), 1 μM (lane 3), 2 μM (lane 4), 4 μM (lane 5), 8 μM (lane 6), and 16 μM (lane 7) StbA_1-75_.

**Figure S6.**
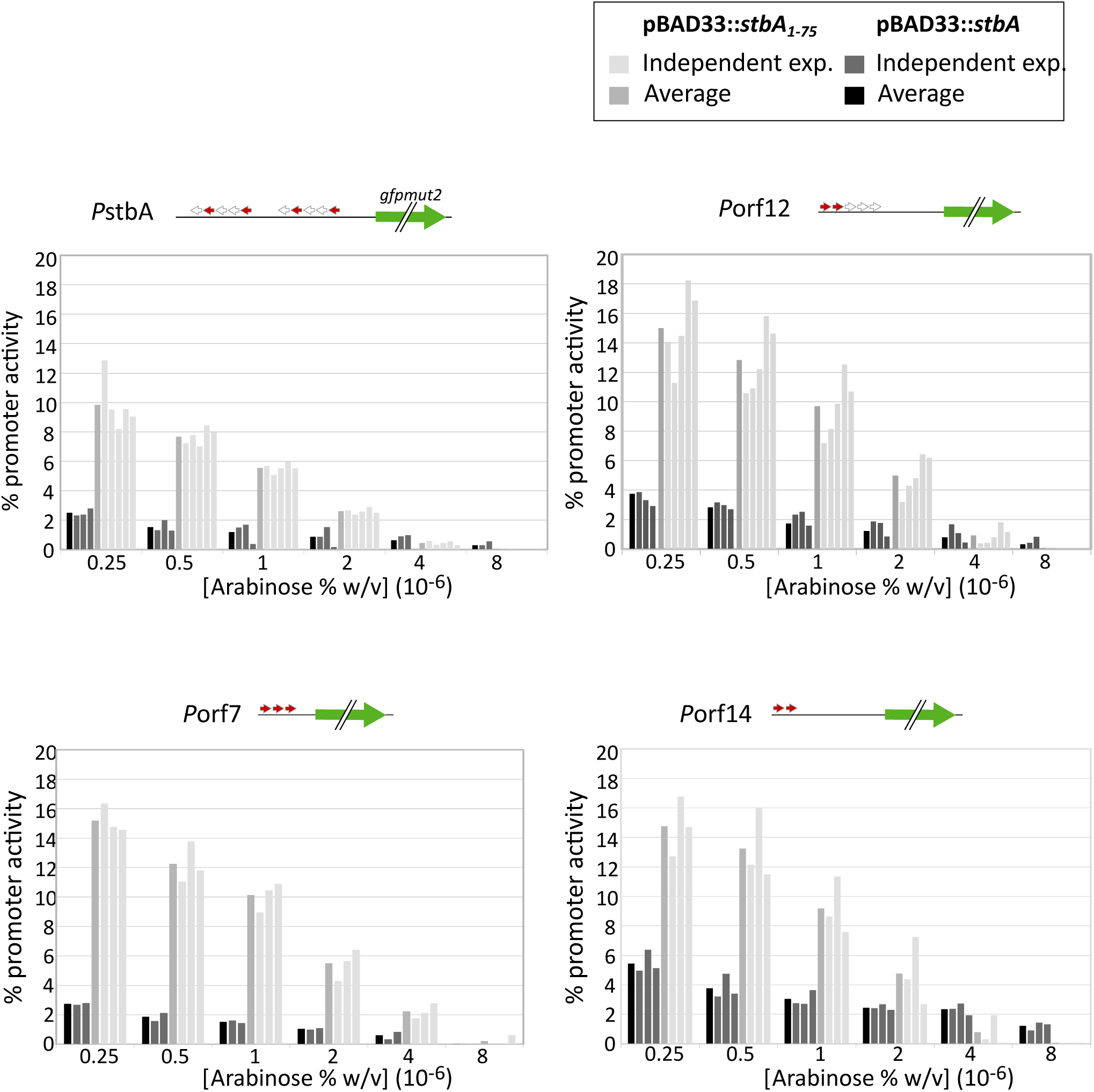
Transcriptional repression activity of StbA and StbA_1-75_ on several *stbDR*-carrying promoters of plasmid R388. The figure shows the percentages of activity of four different promoters carrying *stbDR* sequences in the presence of StbA or StbA_1-75_. StbA or StbA_1-75_ were produced from a co-residing plasmid pBAD33 (pBAD33::*stbA* or pBAD33::*stbA_1-75_*, respectively) and various concentrations of inducer (arabinose) were tested as indicated above the graphs. Percentages of promoter activity are calculated as the ratio between the expression levels (GFP/OD) measured in the presence of pBAD33::*stbA* or pBAD33::*stbA_1-75_* and those measured with the empty vector pBAD33 multiply by 100. Bar charts are described in the legend on the top of the figure. The upper diagrams show the localization and number of *stbDR* in each promoter (gray arrows, consensus in red).

## Notes

### Competing Interest Statement

The authors have declared no competing interest.

